# The optimal docking strength for reversibly tethered kinases

**DOI:** 10.1101/2022.02.07.479365

**Authors:** Mateusz Dyla, Nicolás S. González Foutel, Daniel E. Otzen, Magnus Kjaergaard

**Author notes:** Shared first authors (Mateusz Dyla, Nicolás S. González Foutel). Novozymes A/S, Krogshøjvej 36, 2880 Bagsværd, Denmark (Mateusz Dyla).

## Abstract

Many kinases use reversible docking interactions to augment the specificity of their catalytic domains. Such docking interactions are often structurally independent of the catalytic domain, which allow for flexible combination of modules in evolution and in bioengineering. The affinity of docking interactions spans several orders of magnitude. This led us to ask how the affinity of the docking interaction affects enzymatic activity, and how to pick the optimal interaction module to complement a given substrate. Here, we develop equations that predict the optimal binding strength of a kinase docking interaction and validate it using numerical simulations and steady-state phosphorylation kinetics for tethered protein kinase A. We show that a kinase-substrate pair has an optimum docking strength that depends on their enzymatic constants, the tether architecture, the substrate concentration and the kinetics of the docking interactions. We show that a reversible tether enhances phosphorylation rates most when: I) The docking strength is intermediate, II) the substrate is non-optimal, III) the substrate concentration is low, IV) the docking interaction has rapid exchange kinetics, and V) the tether optimizes the effective concentration of the intra-molecular reaction. This work serves as a framework for interpreting mutations in kinase docking interactions and as a design guide for engineering enzyme scaffolds.

## Introduction

Protein kinases form the backbone of cellular signalling pathways, where they must distinguish their cognate substrates from a wealth of similar protein sequences. The sequence motif surrounding the phosphorylation site only provides part of this specificity as the peptide motifs elucidated *in vitro* do not predict substrate usage *in vivo*. Kinase specificity relies heavily on the local abundance of substrates and enzymes, which means that the same kinase can act in several distinct microenvironments and pathways.^1,2^ The local abundance of kinases relies on either protein interaction domains^3^ or associated anchoring and scaffolding proteins.^4^ These protein interactions can tether kinases to upstream activators or downstream substrates in what is refered to as signalling complexes. When the kinase is tethered to its substrate, phosphorylation occurs inside a complex where the substrate is present at high effective concentration (*C_eff_*). Such tethering can increase the rate of phosphorylation by orders of magnitude^5–8^ and thus increase signaling specificity compared to untethered substrates.

A broad picture of how kinase tethering works has emerged from studies of key model kinases: The active site of the kinase recognise a short linear motif (SLiM) surrounding the phospho-site. A reasonably good motif is required to position the substrate correctly for phosphorylation, but *bona fide* substrates are often far from the consensus sequence. Additionally, many kinases use structurally independent docking interactions, that are not required to position the substrate in the active site, but rather increase the *C_eff_* of the substrate.^6^ Since precise positioning is not required, the docking interaction can take many forms. MAP kinases and cyclin A:cyclin-dependent kinase 2 recognise SLiMs using a binding pocket distant from the catalytic site.^6,9,10^ Other kinases contain dedicated protein interaction domains that typically bind SLiMs such as SH2 or SH3 domains. The kinase and substrate can also be tethered by other proteins, for example in the case of protein kinase A (PKA), the family of more than 50 different A-kinase anchoring proteins (AKAPs).^2^ Many different connections between kinase and substrate can work, as long as the substrate is sterically allowed to bind while the docking interaction is in place.^8^ Therefore, signalling complexes often involve intrinsically disordered regions^11^ that act as flexible chains, and provide the required conformational freedom. The linker architecture also defines the *C_eff_* of the tethered substrate, which regulates the phosphorylation reaction via a Michealis-Menten like dependence on *C_eff_*.^5^ When the linker is longer than needed, the *C_eff_* decreases with linker length following a polymer scaling law, which depends both of the length and chemical composition of the linker.^12^ In total, the specificity enhancement of kinase tethering thus depends on the interplay between what can be considered three interacting modules: The substrate motif, the docking interaction and the linker architecture.

Kinase docking interactions are usually non-convalent and reversible, and their strength can be characterized by rate and equilibrium constants for binding and dissociation (*K_D_*, *k_on_* and *k_off_*). High affinity AKAPs bind with a sub-nanomolar *K_D_*,^13^ whereas short linear motifs typically have *K_D_* values in the micro- to millimolar range.^14^ This translates into complex lifetimes from many minutes to milliseconds. Catalytic efficiency has a non-trivial dependence on docking strength as stronger docking interactions have either been found to decrease^7^ or increase^8^ steady-state phosphorylation by cyclin-dependent kinase, and the same docking interaction affects substrates differently.^15^ This has led to the general conclusion that docking interactions should have a moderate affinity^16^ in analogy with the Sabatier principle in heterogeneous catalysis.^17,18^ The kinetic origins of this principle are illustrated by considering the extremes of strong and weak docking. Weak docking leads to little complex formation, and thus do not enhance phosphorylation rates above the untethered reaction. Strong binding saturates the enzyme with substrate and allows an efficient first cycle of phosphorylation. However, subsequent catalytic cycles are limited by slow product dissociation and thus reduced *k_cat_*. Enzyme docking interactions are prone to this type of product inhibition, as the binding site is structurally independent from the substrate, and does not change in the reaction. Efficient docking interactions must be of intermediate strength to be sufficiently strong to ensure a reasonable bound population and sufficiently weak to allow rapid product dissociation.

What does an optimum at an intermediate binding strength mean in practice? And how does it depend on the properties of the substrate and linker architecture? These are not just questions of fundamental interest, but have practical implications in design of synthetic signalling scaffolds^19^ or interpretation of disease-associate mutations in kinase anchoring proteins. Increased docking strength can either increase or decrease the activity of a pathway, although this cannot be predicted currently even when the molecular details are known. Here, we aim to develop a kinetic framework for predicting how kinase tethering affects steady-state phosphorylation rates depending on the properties of the substrate motif, the docking interaction and the linker architecture. A key goal is to define the optimal docking strength for a system as it allows us to predict whether catalytic efficiency increases or decreases with increased affinity.

Many kinase tethering systems have a modular design with flexibly linked docking interactions. Therefore it is likely possible to develop a kinetic framework that describes many different kinases, or even other classes of similarly tethered enzymes such as phosphatases.^20^ The generality of the approach taken here rests on the assumption that the transient connection between enzyme and substrate only acts as a passive linker that defines *C_eff_*. Sequence-function relationships among IDRs are complex, but the assumption of a passive linker is a good baseline hypothesis for many systems as demonstrated by the successes in describing IDRs using polymer models.^21^

A general kinetic scheme can be studied in a convenient model system such as PKA, which is well-described in terms of structure, mechanism, and kinetics. The catalytic mechanism is similar in other kinases, and is well-captured by standard kinetic descriptions. The key challenge is thus how to represent the structurally diverse connection between kinase and substrates in a way where the properties of the connection can be continuously varied and quantified. We recently developed a model system to study single-turnover reactions of tethered PKA, where the kinase is connected to the substrate via two disordered linkers connected by a heterodimeric coiled-coil (Fig. 1A). The connection can be continuously tuned by changing the lengths of the linkers, which were kept strictly as glycine-serine repeats to avoid sequence-specific effects^5^. Here, we expand this model system by varying the strength of the coiled-coil interaction more than 1.000-fold and investigate how steady-state phosphorylation by reversibly tethered PKA depends on the properties of the substrate, the docking interaction and the linker architecture. Numerical simulations and experiments are used to validate analytically derived rate equations that provide a general framework to predict the effects of enzyme tethering.

**Figure 1:**
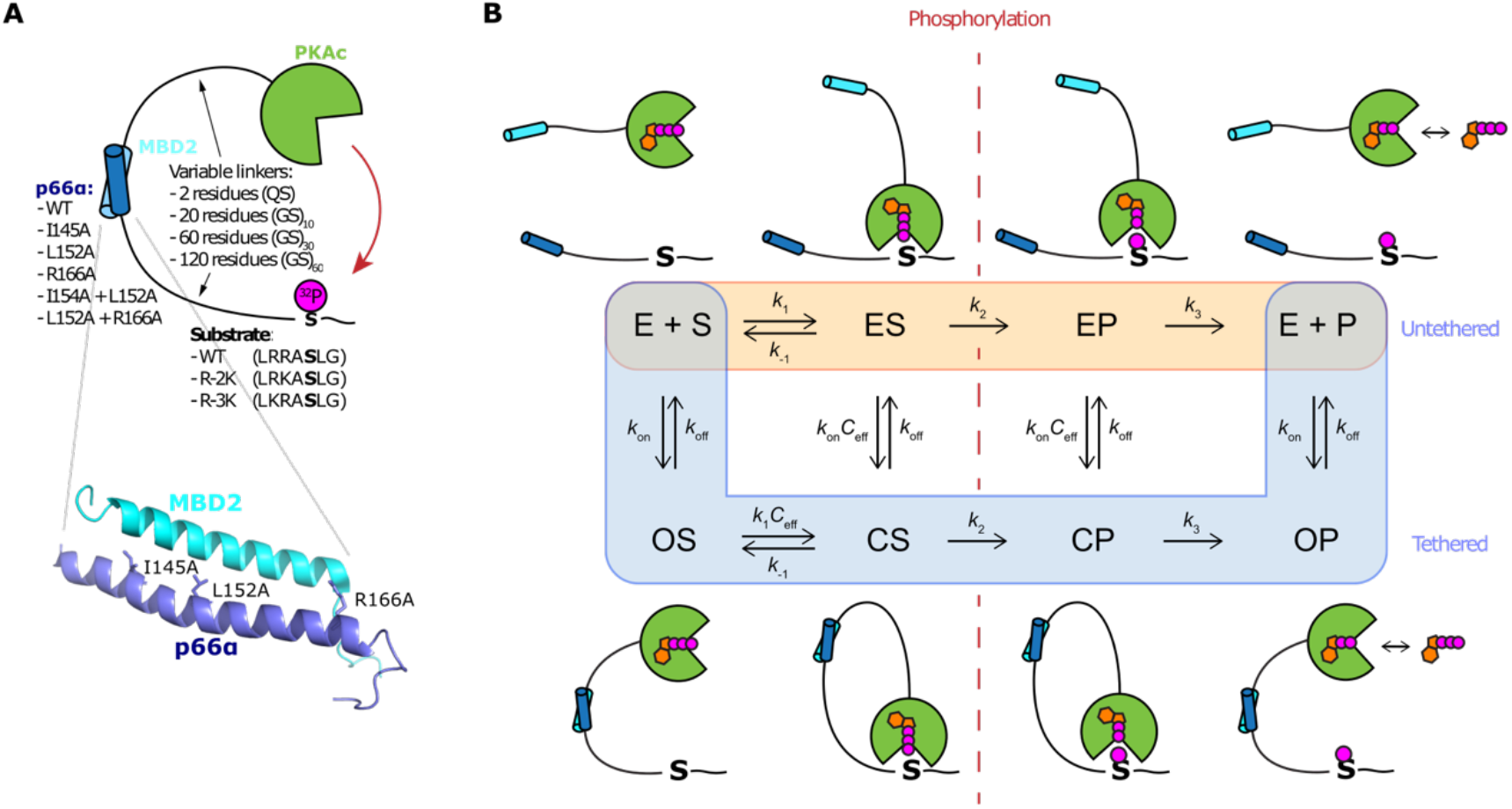
A model system for reversible tethered kinases. **A)** The model system^5^ consists of the catalytic domain from PKA (PKAc) tethered to different substrate motifs via the coiled-coil interaction between p66α and MBD2 (pdb:2L2L)^27^ and disordered GS-linkers. The model system is extended by varying the strength of the docking interaction by mutagenesis of p66α. **B)** Kinetic scheme of the main reaction pathways, where the substrate is either phosphorylated by a tethered or untethered kinase. This scheme omits the ternary complex pathway (Fig. S1) that are kinetically less import. ADP and ATP are represented as orange symbols, but are not explicitly included in the model.

## Materials and Methods

### Preparation ofDNA constructs

Plasmids were prepared by *de novo* synthesis by Genscript and codon optimized for expression in *E. coli*. The coding regions were cloned into pET15b vectors using the NdeI/XhoI sites. The protein sequences and the list of all constructs used are given in the SI.

#### Protein expression and purification

MBD2 peptide containing an N-terminal His-tag and a single C-terminal cysteine (MBD2-Cys) was expressed in BL21(DE3) cells in ZYM-5052 autoinduction medium containing 100 μg/mL ampicillin at 37°C and shaking at 120 RPM for 22 hours. The cells were harvested by centrifugation (15 min, 6000 g), and bacterial pellets were resuspended in a binding buffer (20 mM NaH_2_PO_4_, 0.5 M NaCl, 20 mM imidazole, 0.1 mM TCEP, pH 7.4). To lyse the cells and precipitate folded proteins, the cell suspension was incubated for 25 min at 80°C, while shaking at 400 rpm, before rapidly cooling the cell suspension on ice.^22^ The lysate was centrifuged (15 min, 27000 g) and the supernatant was applied to gravity flow columns packed with Ni-NTA Superflow (QIAGEN). The columns were washed with buffers (20 mM NaH_2_PO_4_, 0.5 M NaCl, 0.1 mM TCEP) containing increasing concentrations of imidazole (40, 60, 80, 100, 200, and 500 mM). An SDS-PAGE gel revealed purified MBD2 peptide in fractions containing 200 and 500 mM imidazole. These fractions were dialyzed against TBS buffer (20 mM Tris-base, 150 mM NaCl, 0.1 mM TCEP, pH 7.6), and subsequently supplemented with TFA to bring the pH to 2, before an additional purification step by reverse-phase chromatography (RPC) in 0.065% TFA buffer, with elution buffer containing also 70% acetonitrile, and using SOURCE 15RPC ST 4.6/100 column (Cytiva). Eluted fractions were lyophilized to dryness.

Protein constructs containing the catalytic domain of PKA linked by (GS)_n_-linker to the MBD2 coiled-coil domain (MBD2-(GS)_n_-PKAc, with n=10, 30 or 60) and an N-terminal His-tag, were expressed in C41(DE3) cells in LB medium with 100 μg/mL ampicillin at 37°C and shaking at 120 RPM. The cultures were induced with 1 mM IPTG at OD_600_ = ~1 and the temperature was decreased to 20 °C for an overnight expression. The cells were harvested by centrifugation (15 min, 6000 × g), bacterial pellets were resuspended in binding buffer (20 mM NaH_2_PO_4_, 0.5 M NaCl, 5 mM imidazole, 0.1 mM TCEP, 0.2 mM PMSF, 50 mg/L of leupeptin, 50 mg/L pepstatin, 50 mg/L chymostatin, pH 7.4) and lysed by sonication (50% duty cycle, maximum power of 70%, sonication time of 5 min). The lysates were centrifuged (20 min, 27000 ×g) and the supernatants were applied to gravity flow columns packed with Ni-NTA Superflow (QIAGEN). The columns were washed with buffers (20 mM NaH_2_PO_4_, 0.5 M NaCl, 0.1 mM TCEP) containing increasing concentrations of imidazole (20, 30 and 40, 150, and 500 mM). SDS-PAGE gels revealed purified PKA samples in fractions containing 40 and 150 mM imidazole. These fractions were upconcentrated (Vivaspin 20 concentrators, 10.000 MWCO) and additionally purified by size-exclusion chromatography (SEC) in TBS buffer (20 mM Tris-base, 150 mM NaCl, 0.1 mM TCEP, pH 7.6) using Superdex 75 Increase column (Cytiva).

Protein constructs containing PKA substrates (WT, R-2K or R-3K) linked to the p66α coiled-coil domain (p66α-QS-substrate, with 6 variants of p66α) and an N-terminal His-tag were expressed in BL21(DE3) cells in Terrific Broth (TB) medium containing 100 μg/mL ampicillin at 37 °C and shaking at 120 RPM. After induction with 1 mM IPTG at OD_600_ = ~0.9, the cultures were allowed to grow for 4 h at 37°C. The cells were harvested by centrifugation (15 min, 6000 ×g), and bacterial pellets were resuspended in binding buffer (20 mM NaH_2_PO_4_, 0.5 M NaCl, 5 mM imidazole, pH 7.4). To lyse the cells and precipitate folded proteins, the cell suspensions were incubated for 25 min at 80 °C, while shaking at 400 rpm, before rapidly cooling the cell suspension on ice. The lysates were centrifuged (15 min, 27000 ×g) and the supernatants were applied to gravity flow columns packed with Ni-NTA Superflow (QIAGEN). The columns were washed with buffers (20 mM NaH_2_PO_4_, 0.5 M NaCl, 0.1 mM TCEP) containing 20 mM of imidazole, and eluted with 500 mM imidazole. Elution fractions were upconcentrated (Vivaspin 20 concentrators, 5000 MWCO) and dialyzed against TBS buffer (20 mM Tris-base, 150 mM NaCl, 0.1 mM TCEP, pH 7.6).

#### Labeling ofMBD2 peptide

For stopped-flow experiments, 100 μM of the MBD2-Cys peptide in TBS buffer was reduced in 1 mM TCEP, and subsequently labelled with 2 mM maleimide-activated Dansyl dye (Sigma-Aldrich) at 4°C overnight, protected from light, and with 15 rpm rolling. The reaction was stopped with 52.7 mM BME, and excess dye was removed by SEC in TBS buffer using Superdex 75 Increase column (Cytiva).

#### Stopped-flow measurements

Binding kinetics of the docking interaction MBD2/p66α were measured using Chirascan spectrometer equipped with a Hg-Xe lamp with a stopped-flow mixing accessory (Applied Photophysics) and monitoring Dansyl fluorescence. Excitation was at 334 nm with a 10 nm bandwidth, and a 495 nm long pass filter was used to monitor the emission. Measurements were done at 30°C in TBS buffer (20 mM Tris-base, 150 mM NaCl, 0.1 mM TCEP, pH 7.6). In order to obtain observed association rate constants (*k*_obs_), MBD2-Cys-Dansyl at a constant concentration of 100 nM was mixed with equal volume of 2.5 - 20 μM of p66α-QS-substrate^WT^. For the lowest affinity p66α^I145A+L152A^-QS-substrate^WT^ mutant, higher concentrations of 12.5 – 100 μM were used to improve the amplitude of the kinetic trace. The traces were fitted with a single exponential association model to obtain *k*_obs_, observed rate constants were plotted versus p66α concentrations and the data were fitted by linear regression to determine the second-order association rate constant (*k*_on_) from the slope of the fitting line. Dissociation rate constants were determined through displacement experiments by first mixing 100 nM of MBD2-Cys-Dansyl with 1 μM of p66α-QS-substrate^WT^, followed by mixing the pre-formed complex with an excess (5 μM) of unlabelled MBD2-Cys that competes for binding to p66α such that MBD2-Cys-Dansyl is mostly unbound at equilibrium. For p66α mutants with low affinity (p66α^I145A+L152A^-QS-substrate^WT^ and p66α^L152A+R166A^-QS-substrate^WT^), higher concentrations were used to improve the amplitude of the kinetic trace (10 μM of p66α-QS-substrate^WT^ and 50 μM of unlabelled MBD2-Cys). The traces were fitted with a single exponential dissociation model, and the observed rate constants are equal to *k*_off_.

### Numerical simulations

Numerical simulations were performed using KinTek Global Kinetic Explorer software^23^ based on a full kinetic model including four reaction pathways: Untethered, tethered (Fig. 1B), and two ternary pathways (Fig. S1). The association rate of the docking interaction was chosen to match WT MBD2:p66a (*k*_on_ = 1.8 · 10^7^ s^-1^ M^-1^), and *k*_off_ was varied from 0.001-1000 s^-1^. The rate of phospho-transfer (*k*2) was set at saturation rate measured by single-turnover kinetics of the tethered reaction (WT: 307 s^-1^, R-2K: 40 s^-1^, R-3K: 14.1 s^-1^).^5^ The association rate constant of all substrates were set at a value typical of short linear motifs (*k*_1_ = 10^7^ s^-1^ M^-1^). The substrate dissociation rate constant (*k*_-1_) was varied such that the resulting *K*_D_ = *k*_-1_/*k*_1_ matches the observed half-saturation point of the tethered phosphorylation reaction (WT: 1420 s^-1^, R-2K: 8370 s^-1^, R-3K: 84000 s^-1^). ADP release is the rate limiting product dissociation step for PKA,^24^ and was estimated to (k_3_ = 42 s^-1^) based on the steady-state kinetics of WT substrate and applied to all substrates. To test the effect of substrate concentration, all three substrates were simulated at concentrations ranging from 0.1 to 100 μM at a fixed *C*_eff_ of 388 μM corresponding to the longest linker used in this study (see below). To test the effect of linker length, *C*_eff_ was varied between 30 and 3000 μM for a fixed concentration of 3 μM of all substrates. To ensure that no more than 10% of the initial substrate was converted to product, the reactions were simulated for reaction time of 300 s with enzyme concentrations at 10 pM. The formation of all phosphorylated reaction species (EP+P+CP+OP+OSP+OPP) in time was subjected to linear regression, and the reaction rate was measured from the slope and converted to specific activity by division with the enzyme concentration.

### Effective concentration calculations

Our tethered kinase systems provide complex linker architectures between kinase and substrate which consist on flexible segments of 20, 60 or 120-GS residues from the enzyme construct, QS plus extra residues from the substrate construct, and a folded coiled-coil p66a-MBD2 heterodimer. To calculate *C_eff_* covering this complexity, a newly developed approach based on conformational ensembles was used.^25^ Briefly, a physically realistic ensemble of the linker was modelled using the “Ensemble Optimization Method” (EOM). As EOM cannot handle heterodimers, the p66a-MBD2 coiled-coil was replaced with a α-helical poly-alanine rod, which has a similar length and produces the same end-to-end distribution as the folded domain.^25^ Flexible segments were simulated as beads using the “Native chain” for an ensemble of 10.000 conformations. The flexible segments were defined from the first residue not defined in the PDB structures of the PKAc (PDB: 2CPK) or the p66a:MBD2 coiled-coil (PDB: 2L2L) and the first residue in the consensus substrate motif. As the flexible linkers are attached to the C-terminus of both coiled-coil segments, the sequence of the linker between the kinase domain and the substrate domain were set in backwards in the EOM simulation. End-to-end distribution and C_eff_ were then calculated for a spacing of 34 Å, which is the distance the linker architecture must span between the active site and the N-terminal attachment site of the PKA catalytic domain (Fig S3).

### Steady-state kinetics

a given PKAc construct linked to the MBD2 coiled-coil domain (MBD2-(GS)_n_-PKAc) was diluted to 10x final concentration in enzyme dilution buffer (50 mM Tris-base, 0.1 mM EGTA, 1 mg/ml BSA, 1 mM TCEP, pH 7.6) and a p66α-QS-substrate was diluted to 10x final concentration in TBS (20 mM Tris-base, 150 mM NaCl, pH 7.6). Steady-state experiments were executed by manual addition of [γ-^32^P]ATP (final concentration 0.1 mM of 100-200 c.p.m. pmol^-1^) into a reaction mix containing 0.2 or 1nM MBD2-(GS)_n_-PKAc and p66α-QS-substrate (final concentration 1, 3 and 10μM) in a reaction buffer (50 mM Tris-base, 0.1 mM EGTA, 10 mM magnesium acetate, 150 mM NaCl, pH 7.6) at 30°C. In evenly distributed time-steps (20, 30 or 60 seconds) 30 μl of the reaction mix were spotted onto a 4 cm^2^ P81 filter disk (Jon Oakhill, St. Vincents Institute of Medical Research) and placed into 75 mM phosphoric acid to quench the reaction. The filters were washed 3 times with 75 mM phosphoric acid, rinsed with acetone, dried, and counted on the ^32^P channel in scintillation counter as c.p.m. Control experiments were performed in similar manner, using TBS buffer instead of substrate in the reaction mix. These procedures were performed in triplicate. Initial velocities were derived from a slope of a linear regression as μM of ^32^P-incorporated to substrate per minute. Phosphorylation rates (*V*_0_/*E*_0_) were then calculated by dividing the slope by the final concentration of PKAc for each experiment (Fig. S4-S9).

## Results

We recently developed a model system to study tethered phosphorylation reactions by single-turnover kinetics.^5^ The system (Fig. 1A) is composed of two parts: The catalytic domain of protein kinase A linked to the MBD2 coiled-coil domain via a flexible Gly-Ser linker of variable length, and a substrate derived from Kemptide^26^ linked to the coiled-coil domain from p66α. The MBD2 and p66α domains associate to form a hetero-dimeric coiled-coil with nanomolar affinity.^27^ The strong docking interactions between the wildtype domains lead to a life-time of the complex which is much longer than the rate of intra-molecular phosphorylation and the reaction is effectively single-turnover at short time scales. Therefore, this system mimicks both high affinity docking interactions and permanent covalent tethering to intramolecular substrates. Weaker docking interactions lead to multiple association-dissociation events within the experimental time frame. Consequently, phosphorylation can occur through several parallel pathways. The most important pathways are the untethered and bimolecularly tethered pathways shown in Fig. 1B, which differ in whether the docking interaction is formed. Additionally, the reaction can occur through a ternary complex, where a tethered kinase phosphorylates a different substrate from the one it is tethered to (Fig. S1).

To adapt this model system to study reversibly tethered kinases, we wanted to vary the strength of the docking interaction, while keeping other parameters constant. We mutated three residues in the coiled-coil interface (Fig. 1A) to alanine to destabilize the complex. We characterized the association and dissociation kinetics of these variants to a fluorescently-labelled MBD2 peptide (Fig. 2, Fig. S2) using stopped-flow fluorimetry. The wild-type interaction has a *K*_D_ of 4.2 nM in agreement with previous studies^27^, and a fast association (*k*_on_ = 1.8’ 10^7^ M^-1^s^-1^) typical of IDP interactions (Table S1). Single point mutations increased the *K*_D_ by up to 61-fold, and their combination into two double mutants further increased *K*_D_ up to ~2400-fold. This reduction of the docking interaction strength occurred mostly through increased *k*_off_ values, which rose by ~500-fold compared to less than 5-fold reduction of the *k_on_* values (Table S1). This was expected as association rates often fall in relatively narrow range in the absence of electrostatic steering or rate-limiting structural changes^28^.

**Figure 2:**
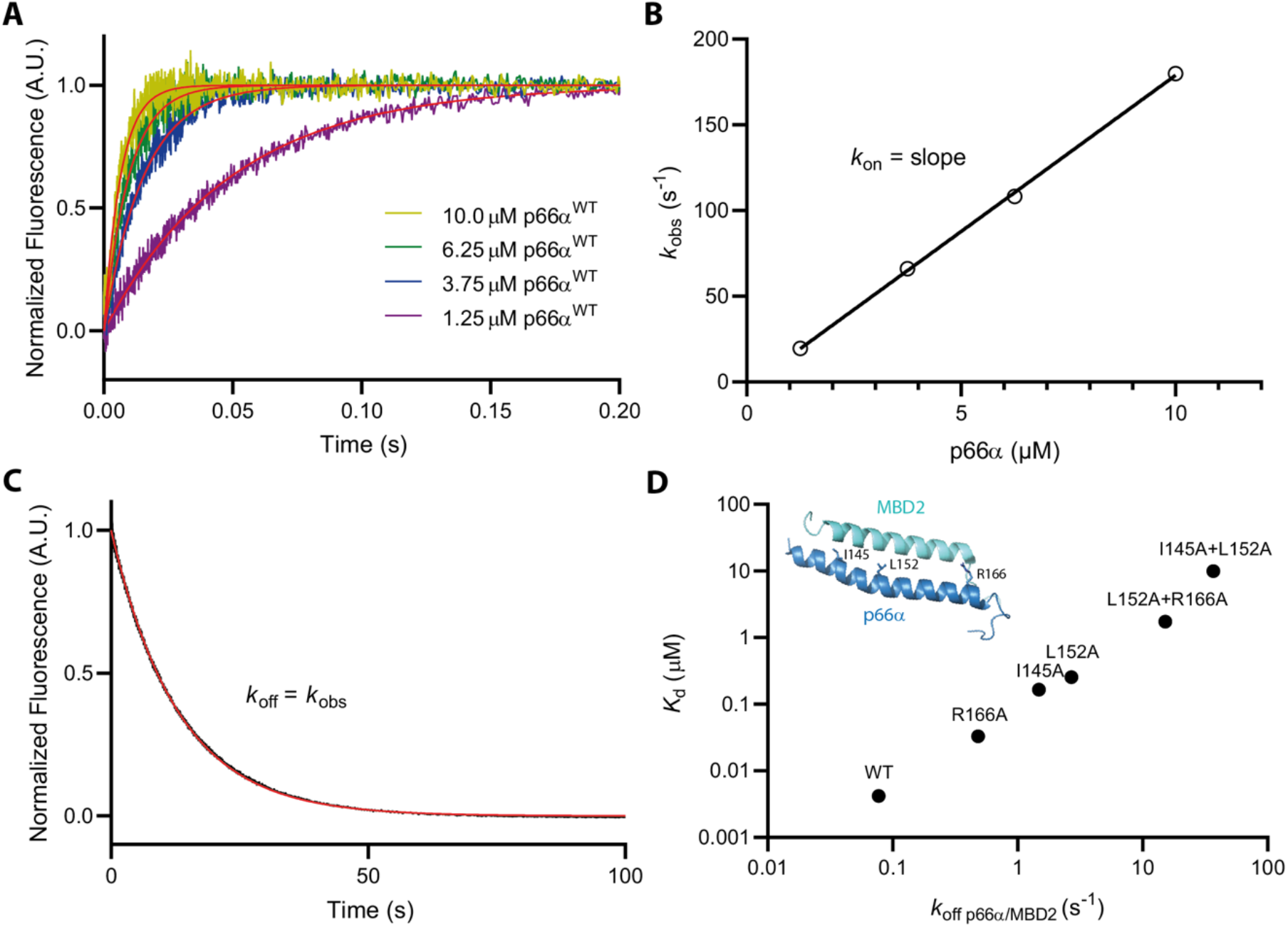
Binding kinetics of variants of the p66α-MBD2 docking interaction by stopped-flow. **A)** Binding of p66α to 50 nM dansyl-labelled MBD2 under pseudo-first order conditions. **B)** The observed rate constant of the fluorescence signal depended linearly on the concentration of p66α with a slope of *k*_on_. **C)** Displacement of 50 nM of dansyl-labelled MBD2 by 2.5 mM of unlabelled MBD2 from a limiting concentration (0.5 mM) of p66α. The observed rate constant is independent of the competitor concentration, which indicates that the observed rate corresponds to a *k*_off_. **D)** The excellent correlation (R = 0.97) between dissociation constants and *k*_off_ in p66α variants shows that the complex is mainly stabilized through a slower dissociation. Error bars representing the standard error from the fit are smaller than the symbols.

We performed numerical simulations to generate theoretical predictions for reversibly tethered kinases, and considered four reaction pathways: Untethered, tethered (both shown in Fig. 1B), and two kinds of ternary complexes where the kinase is attached to a phosphorylated or unphosphorylated substrate (Fig. S1). We simulated phosphorylation of three different variants of the Kemptide substrate: The original Kemptide (WT), which is the optimal substrate for PKA, and two point mutations that reduced *k_cat_/K_M_* by ~10 (R-2K) and ~100-fold (R-3K)^5^, thus comprising a good, an intermediate and a poor substrate. We fixed the *k*_on_ of the docking interaction at the value for wild type MBD-p66a (*k*_on_ = 1.8’ 10^7^ M^-1^s^-1^), which will be similar for many small linear motifs, and varied *k*_off_ to represent docking interactions of various strengths. The effective concentration was determined from an ensemble of 10.000 conforms where MBD2:p66α is simulated as a rigid rod, and disordered segments are treated as a random chain (Fig S3).^25^ For the 120-residue GS-linker this resulted in a calculated *C_eff_* of 388 μM. The numerical simulations predict that steady-state phosphorylation rates have a bell-shaped dependence on the docking strength (*k*_off_) for substrate concentrations from 0.1 to 100 μM (Fig. 3A-C). At high or low docking strengths, the total rate approaches the untethered rate, but reaches an apex at intermediate *k*_off_ values. At higher substrate concentrations, the apex shifts towards weaker docking interactions and the peak broadens. The baseline increases for high substrate concentrations and for better substrates, which suggests a higher flux through the pathways that are independent of the tethering strength, i.e. the untethered and ternary routes.

**Figure 3:**
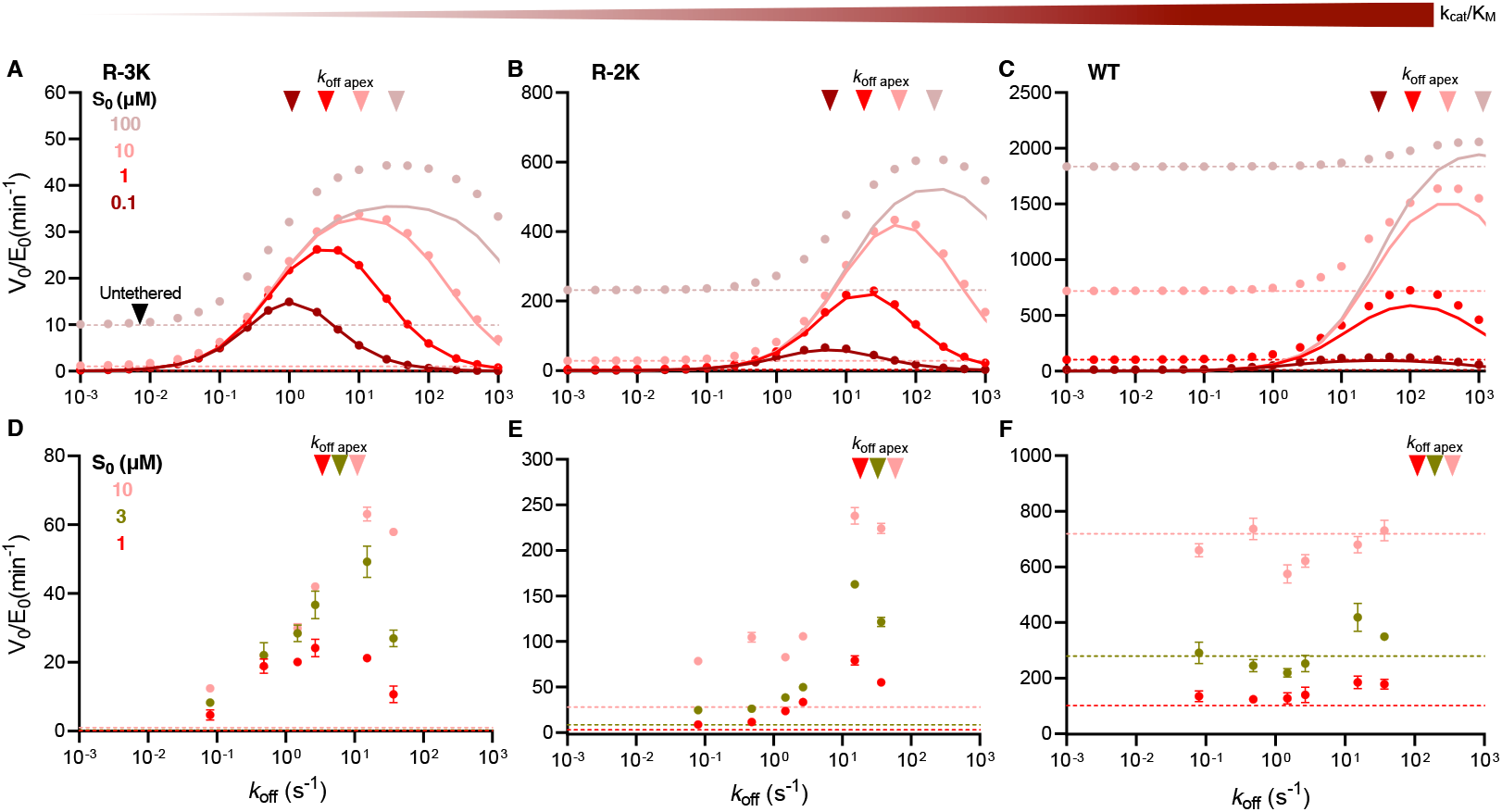
Steady-state phosphorylation rate dependency on substrate concentration from numerical simuations and experiment. **A-C)** Numerical simulation (solid circles) of the steady-state phosphorylation rates for three different PKA substrates: R-3K (A), R-2K (B), and WT (C) at different concentrations. The simulations include the tethered, untethered (Fig. 1B) and ternary reaction paths (Fig. S1). The *k*_on_ was fixed to 1.8 × 10^7^ M^−1^ s^−1^ corresponding to MBD2/p66α, and the *C*_eff_ at 388 μM corresponding to a 120-residues GS-linker. Rate constants for the substrates are based on previous studies.^5^ The dashed lines depict the simulated rates from the untethered reactions with identical parameters (i.e. a parameter that only varies with substrate concentration, not *k*_off_), and the solid lines represent predicted rates from a reaction that only proceeds through the tethered route based on *Eq. I*. Solid triangles indicate the value of *k*_off_apex_ calculated by *Eq. II* (which is based on *Eq. I*) at the simulated conditions. **D-F)** Experimental steady-state phosphorylation rate values for R-3K (D), R-2K (E), and WT (F) substrates at three different substrate concentrations. Error bars represent the standard error of the mean, with n=3. Solid triangles indicate the value of *k_off,apex_* calculated using *Eq. II* for experimental conditions. Unless indicated otherwise, parameters in (D-F) are identical to panels (A-C).

We then sought to develop an analytical solution that could allow us to identify the factors governing the shape and the position of the apex of the bell curves in Fig. 3A-C. It is impractical to consider all four reaction paths in the rate equations, so we inspected the numerical simulations to find reaction paths that could be omitted. The simulations indicated that at low substrate concentrations, the ternary and untethered reaction paths could be ignored. This can be rationalized based on the diffusion-limited access to substrate under these conditions. Therefore, we based our derivation exclusively on the tethered reaction path (Fig. 1B). We derived a kinetic rate equation for product formation under steady-state conditions as a function of the dissociation rate *k*_off_ (SI):

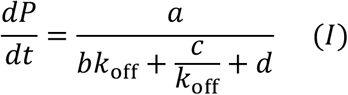

Where a, b, c and d are constants that depend on *S*_0_, *E*_0_, *C*_eff_, *k*_on_, *k*_1_, *k*_-1_, *k*_2_ and *k*_-3_ as defined in the SI. Comparison of predicted kinetics from *Eq. I* to the numerical simulations (Fig. 3A-C) reveals that *Eq. I* describes the reaction kinetics well at low concentrations, but underestimates the phosphorylation rate due to increased flux through the untethered and ternary paths at higher substrate concentrations. However, *Eq. I* still correctly describes the position of the apex seen in the numerical simulations despite significant flux through the untethered path as well as the general behavior of the curve as concentration varies. This observation led us to further calculate the derivative of *Eq. I* relative to *k*_off_, to obtain an expression for docking strength that results in the fastest phosphorylation rate, *k*_off_apex_ (SI):

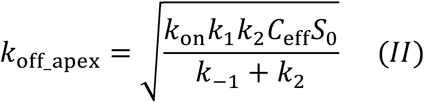

Which for enzymes that follow the Michaelis-Menten reaction scheme (where *k*_2_ = *k*_cat_ and *K*_M_ = (*k*_-1_ + *k*_2_)/ *k*_1_) can be simplified to:

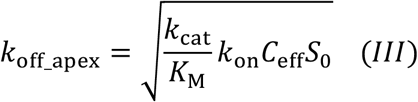

The apex separates two regimes where the strength of the docking interaction affects the catalytic efficiency in opposite directions: At low *k*_off_ values (high docking strength), the docking lifetime is longer than the time required for catalysis. Most docking events lead to phosphorylation and the reaction is limited by how fast the product can be released. Therefore this region is insensitive to the substrate concentration as kinase is saturated with substrate and flux through untethered and ternary paths are neglible. At higher k_off_ values (low docking strength), docking events are shorter than the time required for full phosphorylation, which means that product dissociation is no longer limiting. Instead, the reaction is limited by the fraction of enzymes bound to substrates via the weak docking interaction. *Eq. II* and *III* describe the cross-over between these regimes, and they can be calculated from parameters that can either be measured using standard assays or predicted from physical models.

To test the numerical simulations, we performed a series of steady-state phosphorylation experiments using MBD2-(GS)_120_-PKAc as enzyme and p66α-QS-substrates. For each substrate, we tested our six docking variants at substrate concentrations where the tethered route was predicted to be important corresponding to the middle concentrations used in numerical simulations (1 and 10 μM) and an intermediate concentration of 3 μM (Fig. 3D-F and Fig. S4-S6). Steady-state phosphorylation rates have a bell-shaped dependence on the docking strength for the poor and intermediate (R-3K and R-2K) substrates, whereas no clear apex is seen for the optimal (WT) subtrate. Eq. II shows that the apex for the optimal substrate is expected at higher *k_off_* values than those covered by the range of docking strengths tested. The WT substrate is efficiently phosphorylated regardless of the docking strength with values similar to an untethered reaction (Fig. 3F). A similar situation is seen in the numerical simulations (Fig. 3C), where a small increase is observed on top of a high base line due to a high flux through the non-tethered reactions paths. For the suboptimal substrates, the rates are greatly enhanced and only approach the baseline at high or low *k_off_*. The baseline is defined by the untethered reactions and increase for better substrates and at higher substrate concentrations.

The predicted position of the apex from *Eq. II* vary ~30-fold from the poor to the best substrate (R-3K: *k*_off_apex_ ~3-10 s^-1^ and WT: *k*_off,apex_ ~100-300 s^-1^), and a factor of 3.2 between the highest and lowest measured concentrations (Fig. 3D-F). For both R-3K and R-2K, the predicted apex falls between the two highest observed rates for all concentrations. The change in apex following a 10-fold increase in concentration is less than the spacing between data points and is thus seen as a shift in the relative intensity of the fastest rates. Across this variation of substrate quality and concentration, *Eq. II* and *Eq. III* predict the position of the apex within a factor of ~2-3 despite of the approximations made, thus revealing the predicting capabilities of the analytical solutions.

*Eq. I* also predicts the magnitude of the rate enhancement due to tethering. We found a general agreement between experimental and predicted rate although minor disparities are observed in the amplitudes, for example for R-3K at high concentrations. These differences could be explained by slightly different values of the rate constants that govern substrate binding to the active site (*k*_1_, *k*_-1_) while keeping a constant *k_cat_/K_M_* ratio (Fig. S10). Further assessing of the accuracy and predicting potential of our general equations led us to calculate *Eq. I* using parameters *C_eff_* = 388 μM and [S_0_]= 1, 3 and 10 μM as in our experimental setup (Fig. S11). *Eq. I* appropriately describes the experimental steady-state rate behavior for R-3K and less effectively for WT substrate due to the dominance of the untethered route not considered in the model.

The connection between kinases and their substrates controls the effective concentration of the intra-complex substrate binding interaction and thus impacts the kinetics of the single turnover reaction. However, good substrates are subject to saturation of their single turnover rates in a Michaelis-Menten-like dependence on the effective concentration. This implies that there is a regime where a change in the linker architecture will not affect the rate of phosphorylation. We wanted to test whether transiently tethered kinases are subject to a similar form of *C*_eff_ saturation. In PKA-AKAP signalling complexes, the effective concentration is estimated to vary from the low millimolar to hundreds of micromolar ^29^. We thus performed numerical simulations and compared the three different substrates at a range of effective concentrations that was varied from 30 μM to 3000 μM. We simulated the phosphorylation reaction at 3 μM substrate concentration and fixed association and dissociation rate constants as previously decribed (Fig. 4A-C). Unlike changes in substrate concentration, changes in *C*_eff_ do not influence the flux of the phosphorylation reaction through non-tethered pathways. The relative impact of *C*_eff_ changes are more significant for the poor substrate R-3K: An increase from 300 μM to 3 mM increase phosphorylation rates 5.1-fold at the apex value for R-3K, but only ~1.2-fold for the WT substrate. This is reminiscent of the Michaelis-Menten like dependence observed for tethered single-turnover phosphorylation rates, and occurs for the same reason: For the good WT substrates, the tethered substrate binds fully to the kinase and increases in *C*_eff_ do not increase the fraction of catalytically competent closed complex in contrast to weaker substrates. In addition, as predicted by *Eq. II*, lower effective concentration (longer linkers) pushes the apex towards stronger binding interactions (lower *k*_off_apex_) for all substrates.

**Figure 4:**
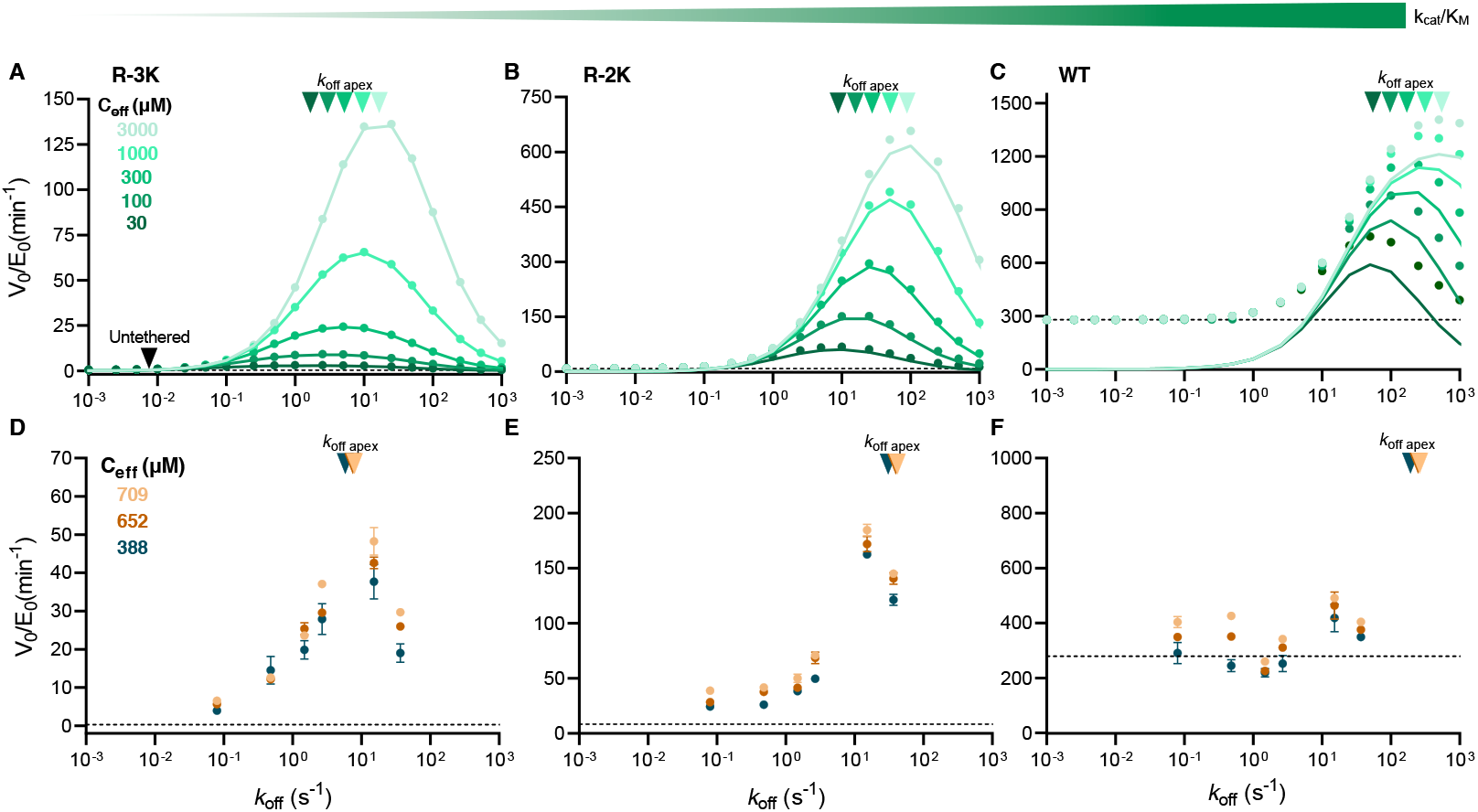
Steady-state phosphorylation rate dependency on effective concentration from numerical simulations and experiments. **A-C)** Numerical simulation (solid circles) of the steady-state phosphorylation rate at different effective concentrations for R-3K (A), R-2K (B) and WT (C) PKA substrates. The simulations include the tethered, untethered and ternary reaction paths. The substrate concentration was set to 3 μM and *k*_on_ and rate constants values for the substrates were fixed as explained in Fig 3. The dashed lines depict the simulated rates from the untethered reactions with identical parameters and the solid lines represent predicted rates from a reaction that only proceeds through the tethered route based on *Eq. I*. Solid triangles indicate the value of *k*_off,apex_ calculated by *Eq. II* with all parameters used in the simulations**. D-F)** Experimental steady-state phosphorylation rate values at three different *C*_eff_= 709, 564 and 388 μM, which correspond to a 20, 60 and 120-residues GS-linker, respectively, for R-3K (D), R-2K (E) and WT (F) substrates. Error bars represent the standard error of the mean, with n= 3. Solid triangles indicate the value of *k_off,apex_* calculated using *Eq. II* for experimental conditions. Unless indicated otherwise, parameters in (D-F) are identical to panels (A-C).

To assess these predictions, we also performed steady-state phosphorylation experiments varying the *C*_eff_. Changes in *C*_eff_ correspond to variations in the connections between enzyme and substrate, which translates into changes in the linker length in our experimental setup. We chose p66α-QS-substrates and three different enzyme constructs: MBD2-(GS)_n_-PKAc with n=10, 30 or 60, and estimated *C*_eff_ values of 709, 652 and 388 μM respectively (Fig. S3). As in the concentration series, we tested docking interaction variants for each substrate quality at a substrate concentration of 3 μM (Fig. 4D-F and Fig. S7-S9). Experiments show that phosphorylation rates vary accompanying changes in effective concentration with a relative effect that depends on subtrate quality and that is more pronounced for *k_off_* values near the apex. However, the observed changes are rather small since the range of *C*_eff_ experimentally tested was limited (~2-fold change in *C*_eff_ from the shortest to the longest linker). For the poor substrate this change in *C*_eff_ is expected to represent ~1.7-fold change in the steady-state rates, whereas for the best substrate only ~1.07-fold change (Fig. S11). The position of the apex (*k_off_apex_*) has a square-root dependence on *C*_eff_ (*Eq. II*), explaining why for our experimental ~2-fold *C*_eff_ change, we observed a small difference in the position in a factor of 1.4. Regardless these small changes, the observed general behavior follows predictions by simulations. For the WT substrate the observed differences in phosphorylation rates are not due to changes in *C*_eff_ but to experimental error, as they assume values near the predicted untethered rate (Fig. 4B).

A given interaction docking strength (*K*_D_) can be achieved by different *k*_on_/*k*_off_ combinations. Using *Eq. I* we calculated the phosphorylation kinetics at association rates ranging from 10^5^ to 10^9^ M^-1^ s^-1^. At comparable *K*_D_ values, fast association-dissociation reactions are more efficient at enhancing the kinase, because the higher dissociation rate allows it to evade the product inhibition that otherwise limits high affinity docking interactions (Fig. S12). This effect is less pronounced for poor substrates as phosphotransfer becomes rate-limiting instead of substrate dissociation. This prediction is difficult to test experimentally as mutations predominantly change the *k_off_* rate, but explains why enzyme targeting often occurs via short linear motifs, which often have higher basal association rates than interactions between two folded domains.^30,31^

## Discussion

We have developed a quantitative model of docking interactions for kinases, which captures the kinase targeting through reversible protein interactions. We compared the model to steady-state phosphorylation in a minimalistic model system, which allowed us to vary docking affinity 2.500-fold, the catalytic efficiency of the substrate up to 100-fold and *C_eff_* ~2-fold. Steady-state kinetics suggest that these parameters affect the phosphorylation reaction by shifting the optimal docking strength and the magnitude of rate enhancement in tethered kinases. The main features of the steady-state kinetics are reproduced by numerical simulations and an equation derived for the tethered reaction path. We suggest that these equations may serve as a first approximation of the effect of mutations and for designing scaffolding interactions.

The equations derived for the tethered reaction path predict how the optimal docking strength depends on the properties of the kinase:substrate complex. Comparisons to steady-state phosphorylation rates show that the equation predicts apex well for the two weaker substrates, whereas the optimal substrate is predicted to have an apex outside the range of docking strengths tested, and is accordingly not observed. When compared to numerical simulations, the analytical equations predict the apex precisely, whereas there are up to 3-fold deviations when compared to experiments. This shows that given precisely defined parameters the equation predicts the apex exactly. Therefore, the deviations are mainly due to the uncertainty in the parameters. However, an error of a factor of 3 in an estimation of the optimal *k*_off_ is in our opinion acceptable for most practical purposes. The total phosphorylation rate also includes the contributions from the untethered reaction paths, which are not considered in *Eq. I–III*. The agreement will become progressively worse as the contribution of this path increases for higher concentrations and better substrates: For R-3K, both the magnitude and curve shape were predicted relatively well, whereas for R-2K, the curve shape is predicted well but the magnitude is off by a factor of 2. For the optimal substrate, the steady-state phosphorylation rate is independent of docking strength throughout the range tested, and the values agree with the calculated untethered rates. Both the numerical simulations and the analytical solutions suggest some enhancement from the tethering in the fast-*k*_off_ regime. This implies that some parameters may not be estimated precisely, likely the rate constants of the substrate binding.

Our model is based on the assumption of modularity, meaning that the substrate-motif, the linker, and the docking motif can be treated as independent modules. This is clearly a simplification, but allowed us to assess the contribution of variable elements in kinase:substrate complex in a general model. Modularity implies structural decoupling between the catalytic domain and the docking interaction, which will typically be the case when they are connected by a disordered linker. Intrinsically disordered regions are common in signalling scaffolds^11^ and kinase anchoring proteins,^29^ which suggests that many docking and catalytic domains will be structurally de-coupled. Furthermore, the architecture of kinases, as well as other enzymes like phosphatases^20,32^, are highly modular with similar catalytic domains coupled to diverse protein interaction domains.^33^ These interactions allow connections to different upstream and downstream partners ultimately making these systems highly evolvable and adaptable^34^. Modularity has been also exploited in biotechnology, bioengineering and pharmacology^35,36,37^ to successfully rewire signaling processes or modify enzymatic activities. This emergent class of engineered multi-domain proteins relies on easy transfer and exchange^34^ of modules, such that design boils down to selecting the right connectors.^38^ For example, synthetic switches with diverse outputs in response to nonphysiological inputs were made by replacing regulatory domains in N-WASP (Wiskott-Aldrich syndrome protein) with heterologous domains and varied linker lengths and substrate-binding affinities.^39^ In other work, modular enzyme scaffold has been used to increase metabolic efficiency.^40^ However, synthetic assemblies do not always work as expected,^41^ suggesting the need for a theoretical framework to guide the design of enzyme tethers.

These and many other examples^34,42^ support this important phenomenom giving solid grounds for our mechanistic framework. However, we cannot predict how more integrated systems, where allosteric coupling between modules or diferent functional units located in a single structure might affect modularity and thus limit the utility of our approach. An example of such system is Protein Phosphatase 1 (PP1), where the RVxF binding motif is located in the same domain as the active site.^20^

Most quantitative studies of enzyme docking interactions considered the catalytic and docking interactions as a single unit. The effect of docking interactions is thus typically to lower *K_M_*, occasionally at the expense of a lowered *k_cat_*. However, tethered enzyme reactions are expected to have non-Michaelis-Menten kinetics as illustrated by numerical simulations (Fig. S13). Furthermore, Michealis-Menten parameters only capture the behavior of a single system, but do not predict the effect of other perturbations than concentration. A theoretical model was proposed describing the effect of the energy of the docking interaction on ERK2 phosphorylation, which predicted the existence of an optimal binding strength although this was not tested experimentally.^43^ This model ignored the untethered reaction path, and is hard to apply in practice as it is formulated in terms of probability functions and inaccessible rates of intra-complex transitions. The competition between the untethered and tethered rate was described in a recent study using a tethered PKA model system.^41^ This study uses a modular description similar to ours, but observes surprisingly low phosphorylation rates. Like our previous work,^5^ this study was aimed at single turnover kinetics, and did thus not consider product dissociation. The model proposed here incooperates ideas from previous treatments into a single framework expressed in terms that can be measured or estimated. The rate enhancement of docking interactions at affinities below the apex is analogous to the decrease in *K_M_*, and the decrease at high affinities is analoguous to decreasing *k_cat_* (Fig. S13). The modular description with focus on the docking interaction allows the effect of perturbations to the system to be predicted, which is attractive for design purposes.

Our kinetic framework can be applied to prediction of the changes associated to mutations in kinases anchoring complexes. Most single nucleotide polymorfisms in docking interactions reduce the affinity. There are many examples from the PKA anchoring proteins (AKAPs): The V282M mutation in AKAP18 reduces the interaction with PKA-RIIα ~9-fold, and impairs cAMP responsive potentiation of L-type Ca^2+^ ion channels^44^. The I646V mutation in the dual-specific D-AKAP2 binds ~3-fold weaker to PKA-RIα, which results in alterations in the subcellular distribution of PKA and cardiac dysfunction.^45^ The S1570L mutation in AKAP9 reduces the interaction with KCNQ1 and consequently its cAMP-induced phosphorylation causing long-QT cardiac syndrome.^46^ Furthermore, in an engineered kinase scaffold composed of tandem PDZ domains kinetic enhancement scaled with PDZ binding affinity.^47^ In all of these cases, weakening of the interaction reduced phosphorylation rates. Examples of mutations that destabilise docking interactions and lead to upregulation of phosphorylation are rare. Phosphorylation by cyclin A-Cdk2 was more efficient when the kinase was docked weakly, rather than more strongly, although the kinetics were not characterized in detail.^7^ In a similar complex, reduction occurs due to a reduced catalytic turnover detected as reduced apparent *k_cat_* values,^15^ which has also been seen for MAP kinases.^48,49^ We lack the complete kinetic parameters to make proper calculations for these systems, but we can qualitatively explain the effect: naturally occurring tethered systems might be optimized around intermediate *K_D_* values. Mutations can reduce catalytic efficiency by either increasing or reducing *k_off_* and *K_D_*, an thus both types of mutations may cause pathological phenotypes. Functional predictions for such mutations do not require accurate calculation of rates, but rather only to know on which side of the apex a given system is. Therefore *Eq. I–III* may be applicable even when some rates are only crudely estimated.

Kinase signaling is rarely optimized for catalytic efficiency, but rather for switchability. In terms of tethering interactions, this can be understood as the relative increase in phosphorylation rates in the presence vs. in the absence of a tethering interaction (Fig. 5A). Analoguously, it can be understood as the ability for a kinase to distinguish the cognate tethered substrate from a similar non-cognate substrate. We thus compared how the kinetic discrimination between an identical tethered and untethered substrate depends on the system parameters (Fig. 5B-D), and formulated five general principles of for when tethered systems maximize enhancement: (I) Enzyme and substrate should interact with intermediate docking strength (typically, *K_D_* in μM-range); (II) Docking interaction should have fast exchange kinetics (high *k_on_/k_off_*) (Fig. S12); (III) The substrate should have low affinity for the enzyme (Fig 5B); (IV) The substrate concentrations should be low (Fig 5C); and (V) The connection should be optimized to enforce high effective concentrations (Fig 5D). As tethered systems with modular design are not only restricted to kinases (see above), other enzymes can be studied through the same framework and these rules can be applied as “design principles” for novel or improved scaffolding interactions. For example, for proximity-based drugs like PROTACs, it has been recognized that a major obstacle is to generate a design with predictable effects.^36^ Effectiveness of PROTACs lies on the likelihood to form stable ternary complexes as well as their final concentration after administration.^36^ In this sense, our approach could be a powerful toolkit for rationally design interactions mediated by scaffolds, optimize drug design/administration as well as predict outcomes of therapeutic treatments.

**Figure 5:**
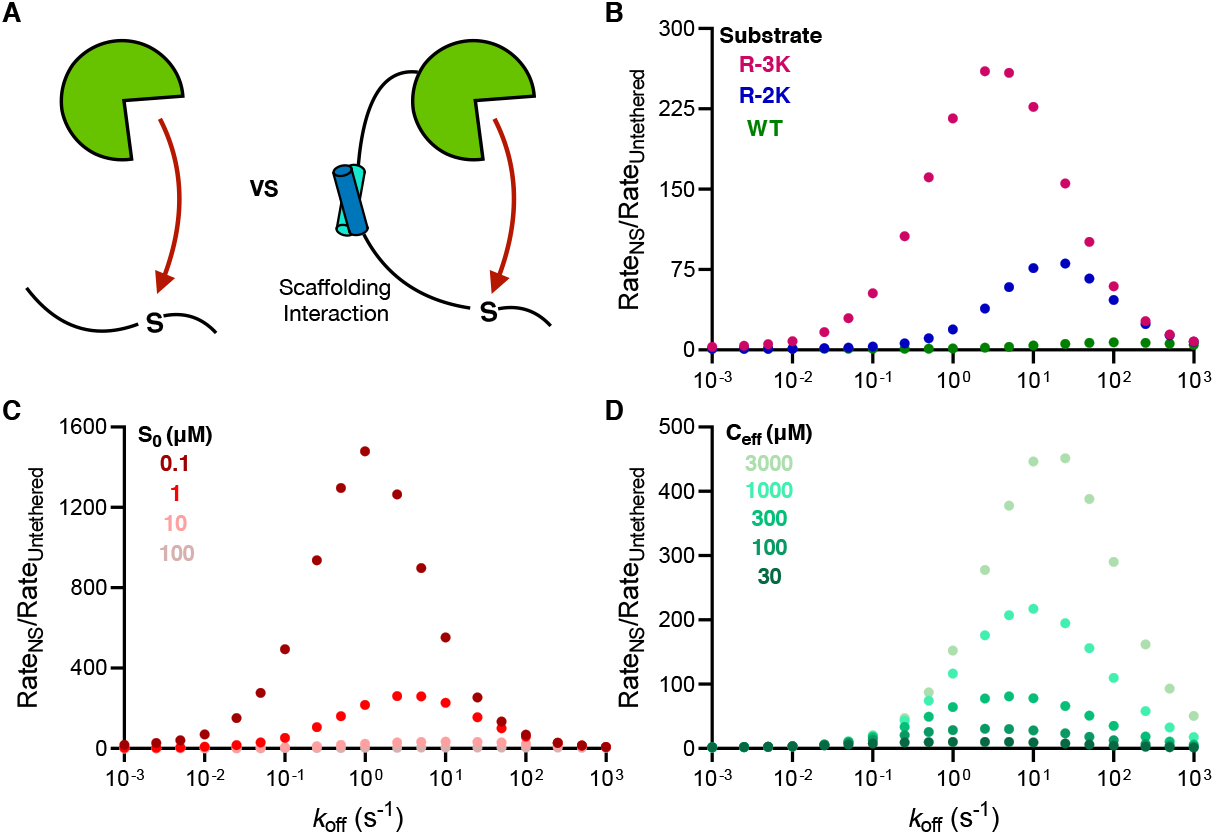
Kinetic discrimination between tethered and untethered substrates. **A)** Schematic representation of the untethered vs reversibly tethered kinase model, where scaffolding interaction results in kinetic rate enhancement. **B-D)** Comparision of the ratio of a tethered reactions predicted by Eq. 1 to the corresponding untethered reaction predicted by the Michaelis-Menten equation. Unless otherwise mentioned, the substrate is R-3K, [S] = 1 μM and *C*_eff_ = 388 μM. The largest kinetic boost is observed **B)** for poor substrates, **C)** at low substrate concentrations, and **D)** at high effective concentrations of substrate.

## Acknowledgments

This work was supported by grants to M.K. from the “Young Investigator Program” of the Villum Foundation, PROMEMO – Center for Proteins in Memory, a Center of Excellence funded by the Danish National Research Foundation (grant number DNRF133) and the Novo Nordisk Foundation (NNF200C0063808). D.E.O. is grateful for support from the Novo Nordisk Foundation (NNF170C0028806) and the Lundbeck Foundation (R276-2018-671).

