## Supplemental information for "The optimal docking strength for reversibly tethered kinases"

[$] Shared first authors

### Present address: Novozymes A/S, Krogshøjvej 36, 2880 Bagsværd, Denmark

### Contents:

p. 3 Overview of protein sequences and constructs used

#### Supplementary figures:

p. 4 **Fig. S1:** Schematic of additional kinetic pathways including ternary complexes

p. 5 **Fig. S2**: Stopped-flow data of binding and dissociation kinetics of p66a variants with MBD2

p. 6 **Fig. S3**: Estimation of effective concentrations of substrate for three different linker architectures.

p. 7-12 **Fig. S4-9:** Raw data for measurement of steady-state phosphorylation rates

p.13 **Fig. S10:** Steady-state phosphorylation rate dependence on substrate binding kinetics

p.14 **Fig. S11:** Comparison of rates from experiment and predicted from analytical solutions

p.15 **Fig. S12:** Phosphorylation rate dependence on the kinetics of the docking interaction

p.16 **Fig. S13:** Phosphorylation rate enhancement dependence on substrate concentration.

#### Supplementary tables:

p.17 **Table S1:** Rate constants from stopped-flow experiments.

#### Supplementary text:

p.18 Derivation of equations describing tethered phosphorylation

#### Protein sequences

**Color coding:**

- 6xHis-tag
- Thrombin cleavage sequence
- MBD2 dimerization domain
- p66α dimerization domain, residues **I145**, **L152**, and **R166** that can be mutated to alanines are highlighted
- NheI restriction site. DNA: GCTAGC. Protein: AS -
- $(GS)_n$ – variable-length GS linker; n = 10, 30 or 60
- KpnI restriction site. DNA: GGTACC. Protein : GT
- PKA substrate motif, catalytic serine is shown in **bold**

**MBD2-$(GS)_n$-PKAc**

MGSSHHHHHHSSGLVPRGSHMVTDEDIRKQEERVQQVRKKLEEALMADAS$(GS)_n$GTGNAAAAKKGSEQESVKEFL
AKAKEDFLKKWETPSQNTAQLDQFDRIKTLGTGSFGRVMLVKHKESGNHYAMKILDKQKVVKLKQIEHTLNEKRIL
QAVNFPFLVKLEFSFKDNSNLYMVMEYVAGGEMFSHLRRIGRFSEPHARFYAAQIVLTFEYLHSLDLIYRDLKPEN
LLIDQQGYIQVTDFGFAKRVKGRTWTLCGTPEYLAPEIILSKGYNKAVDWWALGVLIYEMAAGYPPFFADQPIQIY
EKIVSGKVRFPSHFSSDLKDLLRNLLQVDLTKRFGNLKNGVNDIKNHKWFATTDWIAIYQRKVEAPFIPKFKGPGD
TSNFDDYEEEEIRVSINEKCGKEFTEF

**MBD2-Cys**

MGSSHHHHHHSSGLVPRGSHMVTDEDIRKQEERVQQVRKKLEEALMADASCGSGSGSGSGSY

**p66α-QS-substrate$^{WT}$**

MGSSHHHHHHSSGLVPRGSHMTSPEERERM**I**KQLKEE**L**RLEEAKLVLLKKL**R**QSQIQKEATAQKASQSGTPGSGSG
SGSLRRA**S**LGGGGGY

**p66α-QS-substrate$^{R-2K}$**

MGSSHHHHHHSSGLVPRGSHMTSPEERERM**I**KQLKEE**L**RLEEAKLVLLKKL**R**QSQIQKEATAQKASQSGTPGSGSG
SGSLRKA**S**LGGGGGY

**p66α-QS-substrate$^{R-3K}$**

MGSSHHHHHHSSGLVPRGSHMTSPEERERM**I**KQLKEE**L**RLEEAKLVLLKKL**R**QSQIQKEATAQKASQSGTPGSGSG
SGSLKRA**S**LGGGGGY

#### Table of substrate and enzyme constructs used in this work:

| Substrate | Substrate | Substrate | Enzyme |
|---|---|---|---|
| p66α$^{WT}$-QS-WT | p66α$^{WT}$-QS-R-2K | p66α$^{WT}$-QS- R-3K | MBD2-$(GS)_{10}$-PKAc |
| p66α$^{R166A}$-QS-WT | p66α$^{R166A}$-QS- R-2K | p66α$^{R166A}$-QS- R-3K | MBD2-$(GS)_{30}$-PKAc |
| p66α$^{L152A}$-QS-WT | p66α$^{L152A}$-QS- R-2K | p66α$^{L152A}$-QS- R-3K | MBD2-$(GS)_{60}$-PKAc |
| p66α$^{I145A}$-QS-WT | p66α$^{I145A}$-QS- R-2K | p66α$^{I145A}$-QS- R-3K | |
| p66α$^{L152A/R166A}$-QS-WT | p66α$^{L152A/R166A}$-QS- R-2K | p66α$^{L152A/R166A}$-QS-R-3K | |
| p66α$^{I145A/L152A}$-QS-WT | p66α$^{I145A/L152A}$-QS- R-2K | p66α$^{I145A/L152A}$-QS-R-3K | |

#### Supplementary figures

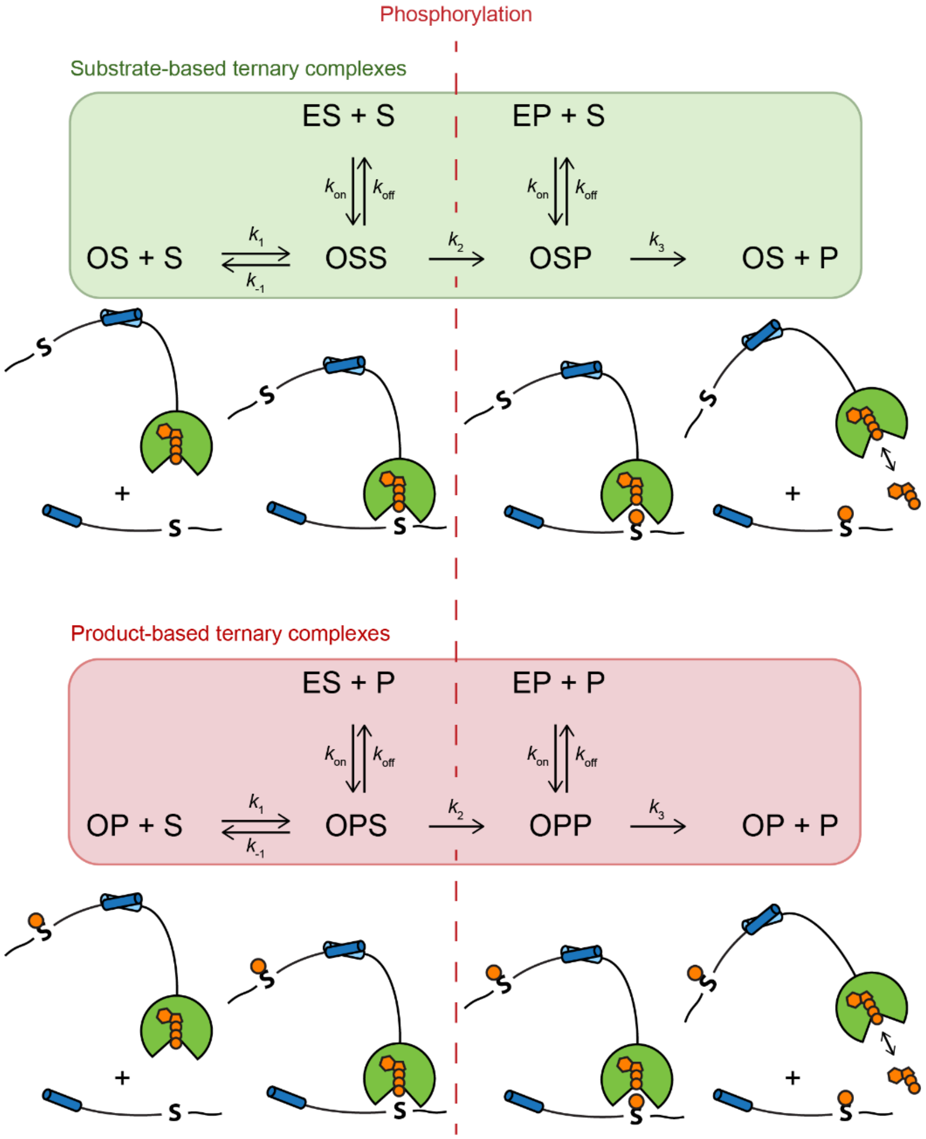


**Figure S1. Additional kinetic pathways composed of substrate-based (upper panel) or product-based (lower panel) ternary complexes.** Ternary complexes will mostly contribute to the reaction scheme at high substrate concentrations and are omitted in the analytical solutions, but are included in the numerical simulations.

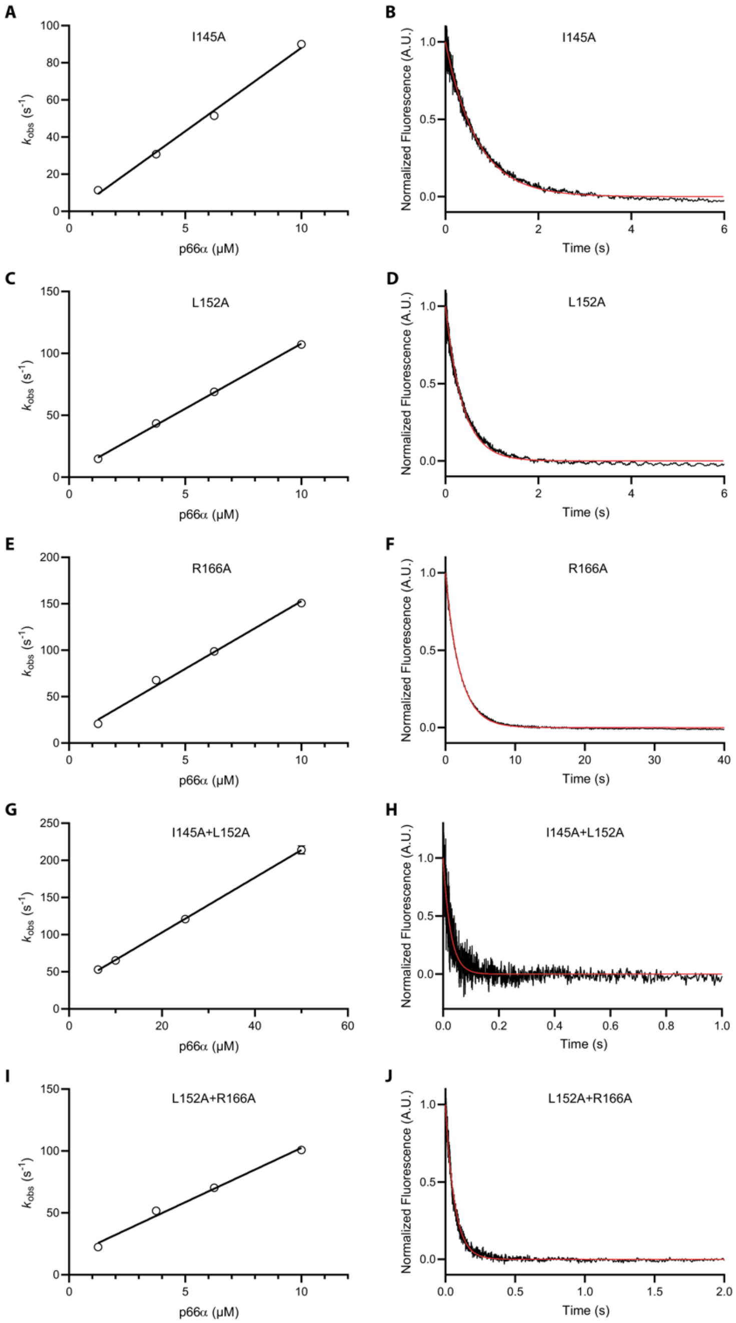


**Figure S2: Stopped-flow data describing the binding and dissociation kinetics of p66α variants with MBD2.** (A, C, E, G, I) The observed rate of change of the fluorescence emision scales linearly with p66α concentration under pseudo-first order conditions and allows the bimolecular association rate constant ($k_{on}$) to be extracted as the slope. The concentration of fluorescently-labelled MBD2 was 50 nM. Error bars representing the standard error from the fit are in most cases smaller than the symbols. (B, D, F, H, J) Competetive displacement of 50 nM of fluorescently-labelled MBD2 from limiting amounts of p66α (0.5 μM for single-mutants (B, D, F) and 5 μM for double mutants (H, J)) by excess unlabelled MBD2. The concentration of unlabelled MBD2 was 2.5 μM for single-mutants (B, D, F) and 25 μM for double mutants (H, J).

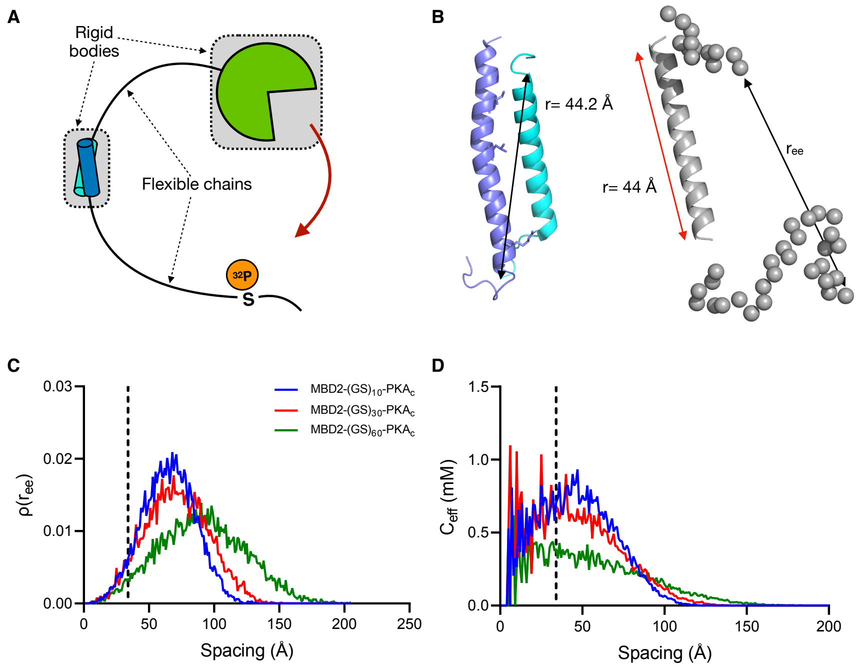


**Figure S3: Estimation of effective concentrations of substrate for three different linker architectures.** (A) The linker architecture connecting the kinase and its substrate can be decomposed into rigid bodies and flexible chains for the purpose of modelling. (B) An α-helical rod was used to replace the p66α:MBD2 coiled-coil (PDB: 2L2L) in the EOM modelling, as described previously[1]. Flexible segments attached to the rod were simulated as beads for an ensemble of 10.000 conformations and one of them is here represented. (C) End-to-end distance distribution and (D) predicted $C_{eff}$ of the three different linker architectures from the ensemble. The spacing corresponding to the distance between active site and N-terminus of PKA is indicated as a dashed line. $C_{eff}$ was 709, 651 and 388 μM for linkers with a 20, 60 and 120 GS-residues respectively between the kinase and the coiled-coil, for a spacing of 34 Å corresponding to the distance between the active site and attachement site on the kinase.

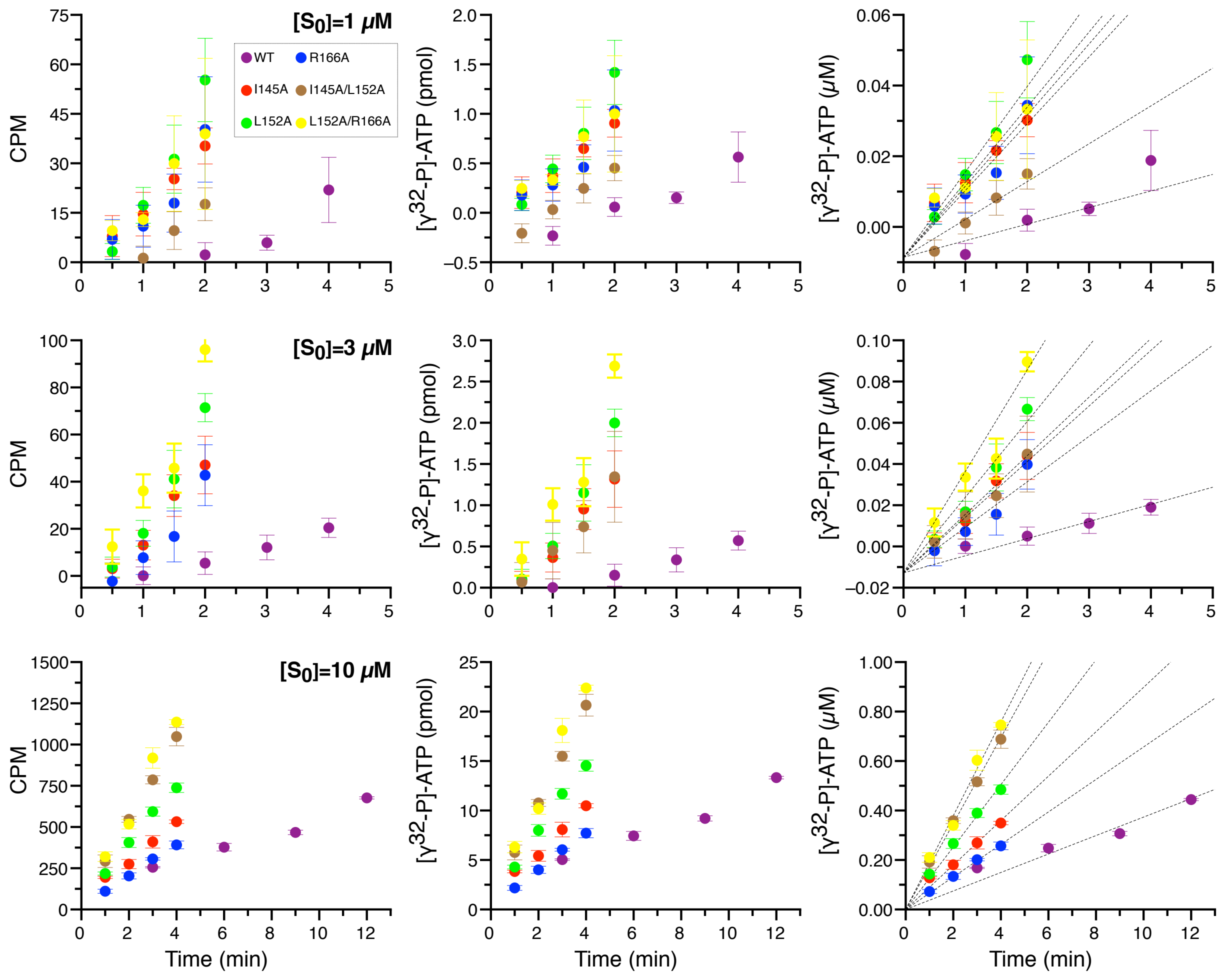


**Figure S4: Measurement of steady-state phosphorylation rates in vitro of p66α-QS-R-3K substrate at three concentrations.** Corrected counts per minute (CPM) from $^{32}P$-incorporated to substrate, obtained at different time-steps for each docking interaction variant. Each point was measured by triplicate (Left plot). Pmol of [$\gamma^{32}P$]-ATP incorporated per minute, obtained after dividing CPM by the slope of the [$\gamma^{32}P$]-ATP standard curve (Middle plot). Initial velocities were obtained from the slope of the linear regression of μM of $^{32}P$-incorporated to substrate per minute (first-order conditions). Error bars represent the standard error of the mean, with n= 3.

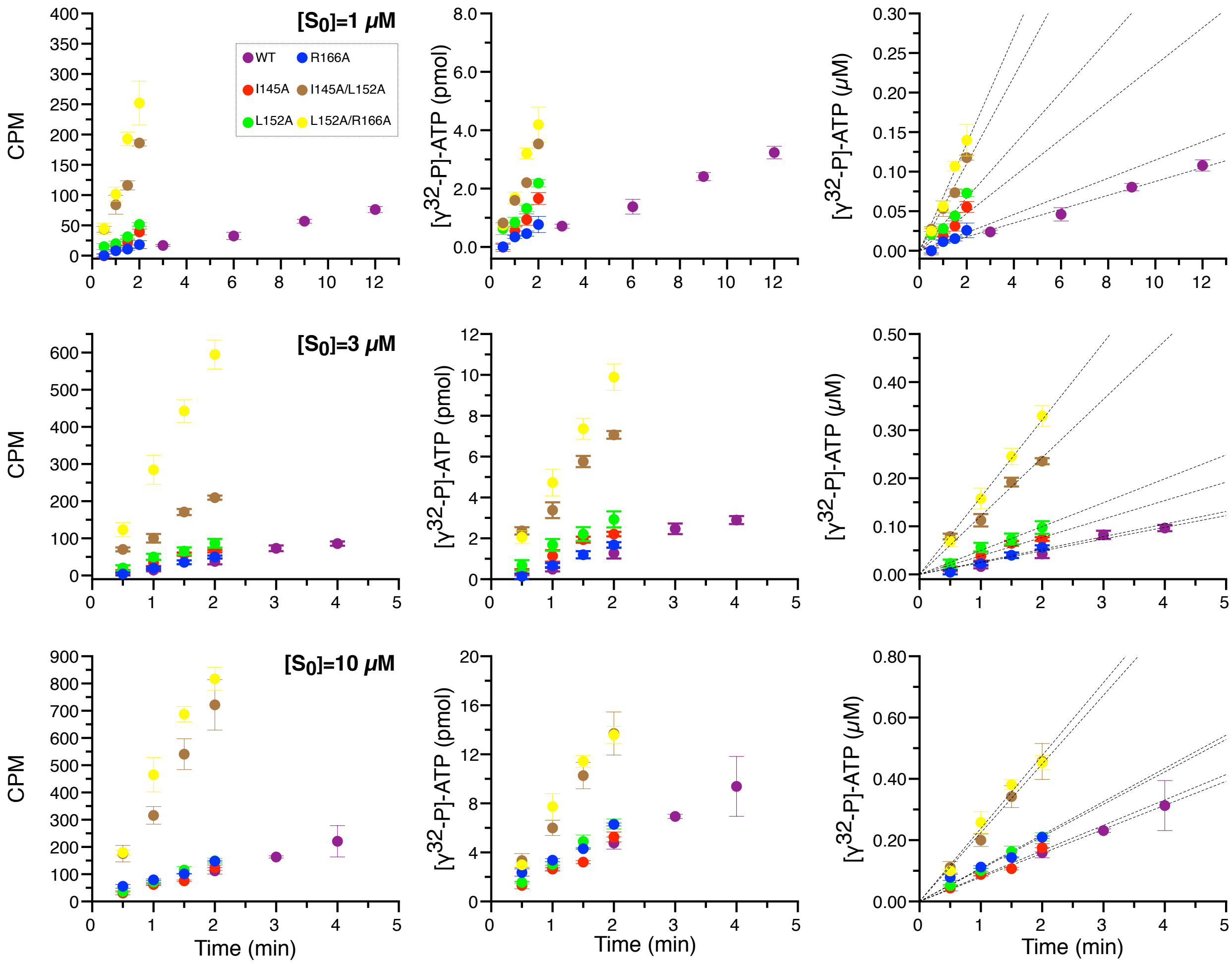


**Figure S5: Measurement of steady-state phosphorylation rates in vitro of p66α-QS-R-2K substrate at three concentrations.** Corrected counts per minute (CPM) from $^{32}$P-incorporated to substrate, obtained at different time-steps for each docking interaction variant. Each point was measured by triplicate (Left plot). Pmol of [$\gamma^{32}$P]-ATP incorporated per minute, obtained after dividing CPM by the slope of the [$\gamma^{32}$P]-ATP standard curve (Middle plot). Initial velocities were obtained from the slope of the linear regression of μM of $^{32}$P-incorporated to substrate per minute (first-order conditions). Error bars represent the standard error of the mean, with n= 3.

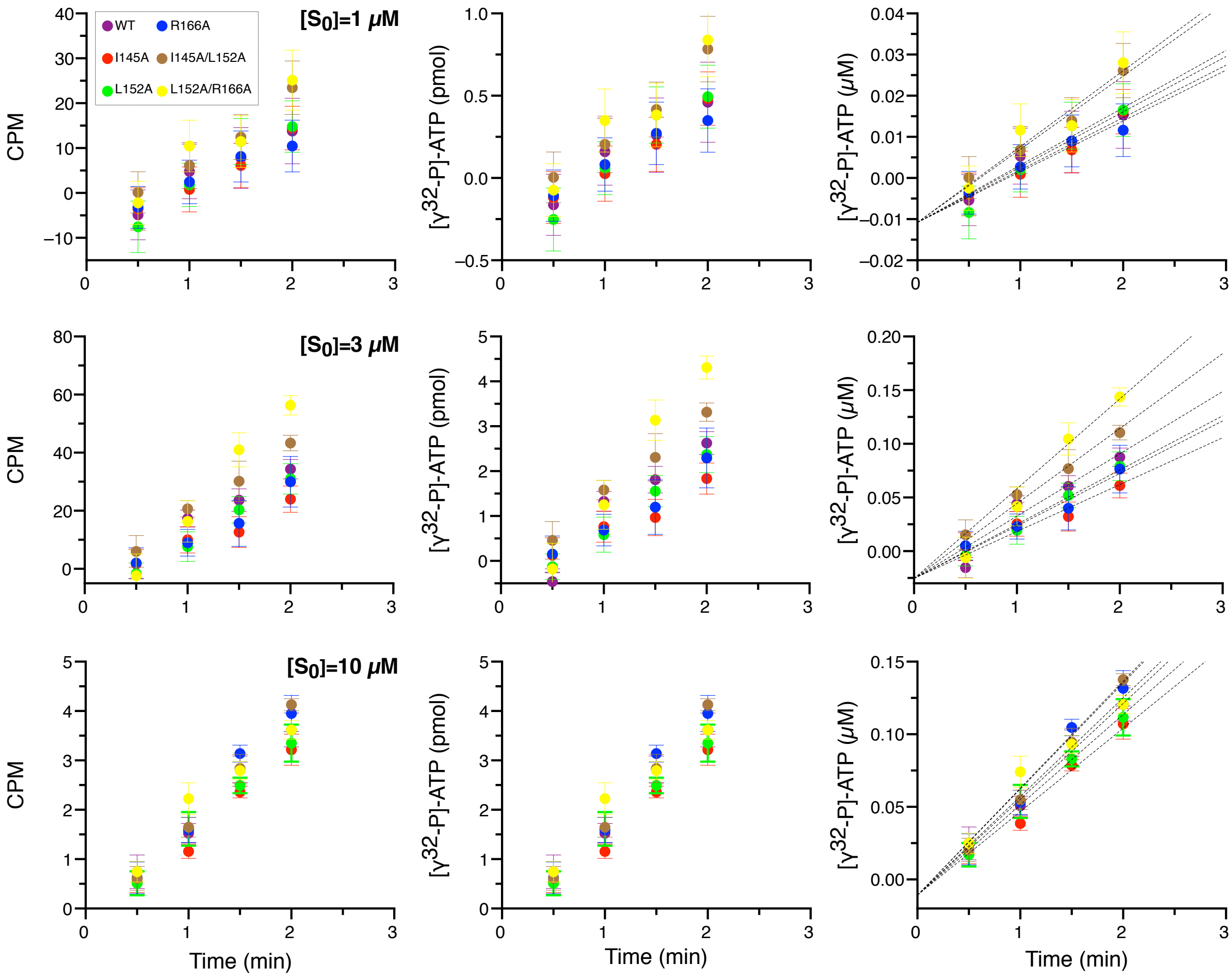


**Figure S6: Measurement of steady-state phosphorylation rates in vitro of p66α-QS-WT substrate at three concentrations.** Corrected counts per minute (CPM) from $^{32}$P-incorporated to substrate, obtained at different time-steps for each docking interaction variant. Each point was measured by triplicate (Left plot). Pmol of [$\gamma^{32}$P]-ATP incorporated per minute, obtained after dividing CPM by the slope of the [$\gamma^{32}$P]-ATP standard curve (Middle plot). Initial velocities were obtained from the slope of the linear regression of μM of $^{32}$P-incorporated to substrate per minute (first-order conditions). Error bars represent the standard error of the mean, with n= 3.

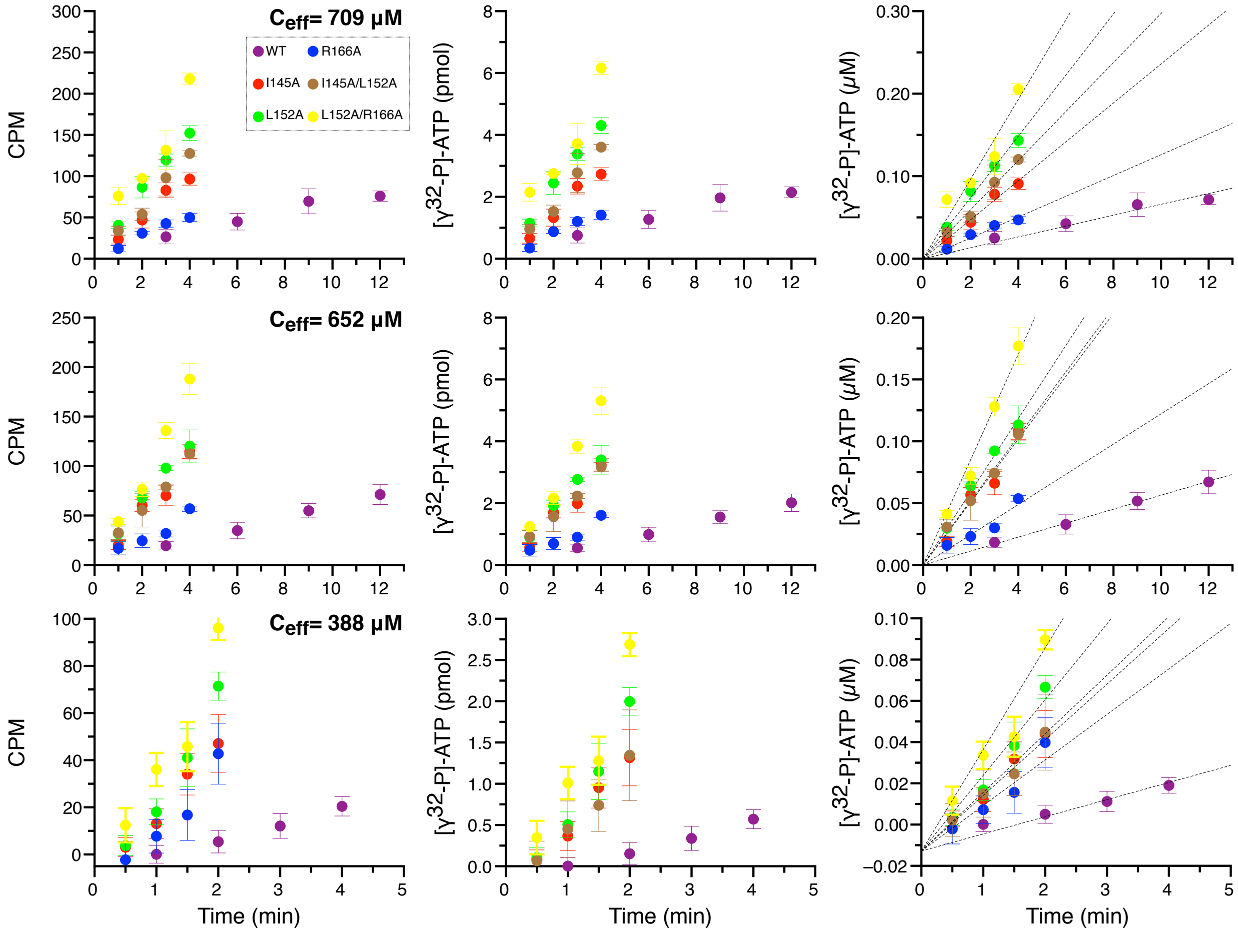


**Figure S7: Measurement of steady-state phosphorylation rates in vitro of p66α-QS-R-3K substrate for three different effective concentracions.** These measurements were performed at 3 μM substrate concentration and using enzyme constructs: MBD2-(GS)$_{10}$-PKAc ($C_{eff}$ =709 μM), MBD2-(GS)$_{30}$-PKAc ($C_{eff}$ =652 μM) and MBD2-(GS)$_{60}$-PKAc ($C_{eff}$ =388 μM). Corrected counts per minute (CPM) from $^{32}$P-incorporated to substrate, obtained at different time-steps for each docking interaction variant. Each point was measured by triplicate (Left plot). Pmol of [γ$^{32}$P]-ATP incorporated per minute, obtained after dividing CPM by the slope of the [γ$^{32}$P]-ATP standard curve (Middle plot). Initial velocities were obtained from the slope of the linear regression of μM of $^{32}$P-incorporated to substrate per minute (first-order conditions). Error bars represent the standard error of the mean, with n=3.

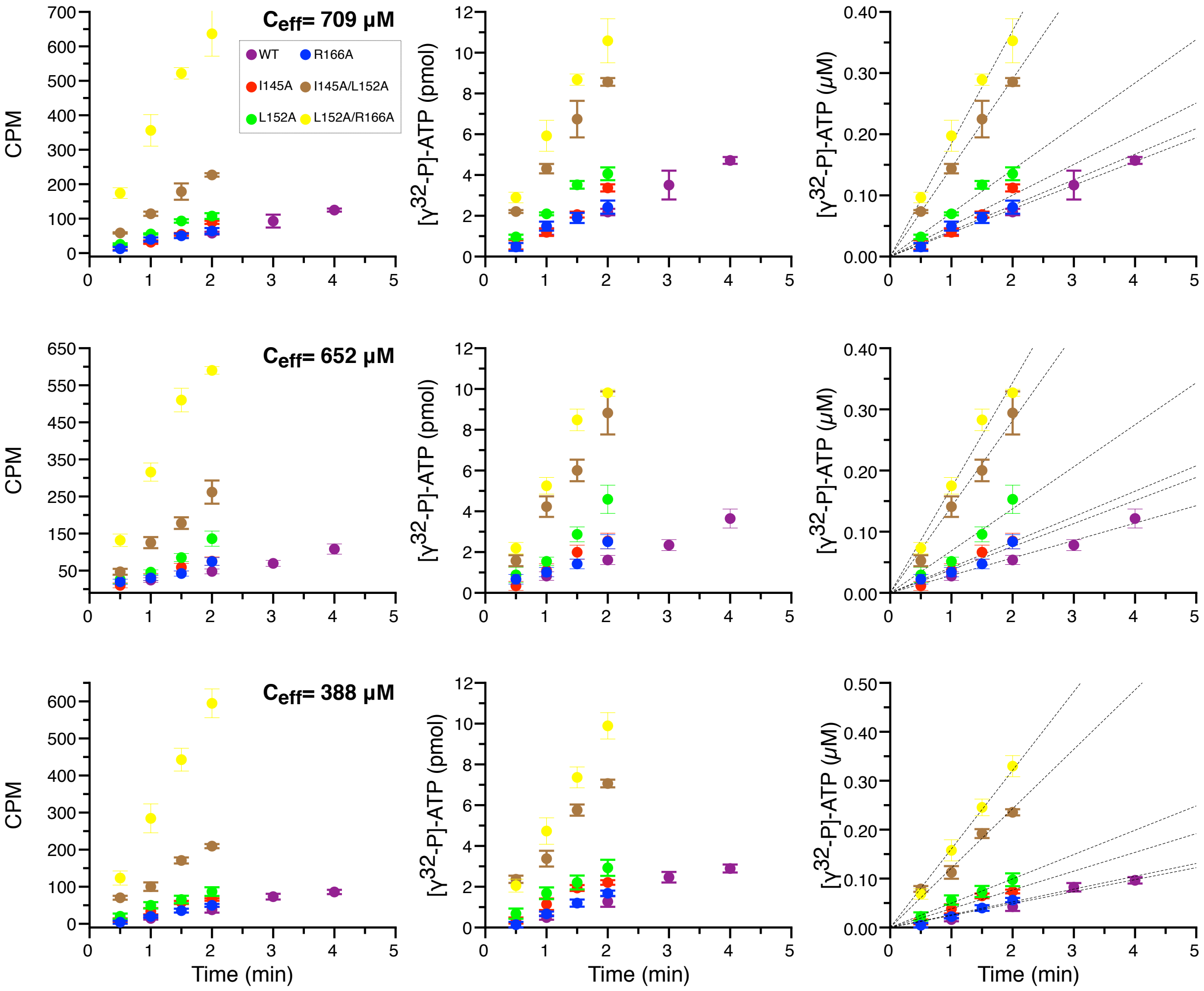


**Figure S8: Measurement of steady-state phosphorylation rates in vitro of p66α-QS-R-2K substrate for three different effective concentracions.** These measurements were performed at 3 µM substrate concentration and using as enzyme constructs: MBD2-$(GS)_{10}$-PKAc ($C_{\mathrm{eff}}$ =709 µM), MBD2-$(GS)_{30}$-PKAc ($C_{\mathrm{eff}}$ =652 µM) and MBD2-$(GS)_{60}$-PKAc ($C_{\mathrm{eff}}$ =388 µM). Corrected counts per minute (CPM) from $^{32}$P-incorporated to substrate, obtained at different time-steps for each docking interaction variant. Each point was measured by triplicate (Left plot). Pmol of [$\gamma^{32}$P]-ATP incorporated per minute, obtained after dividing CPM by the slope of the [$\gamma^{32}$P]-ATP standard curve (Middle plot). Initial velocities were obtained from the slope of the linear regression of µM of $^{32}$P-incorporated to substrate per minute (first-order conditions). Error bars represent the standard error of the mean, with n= 3.

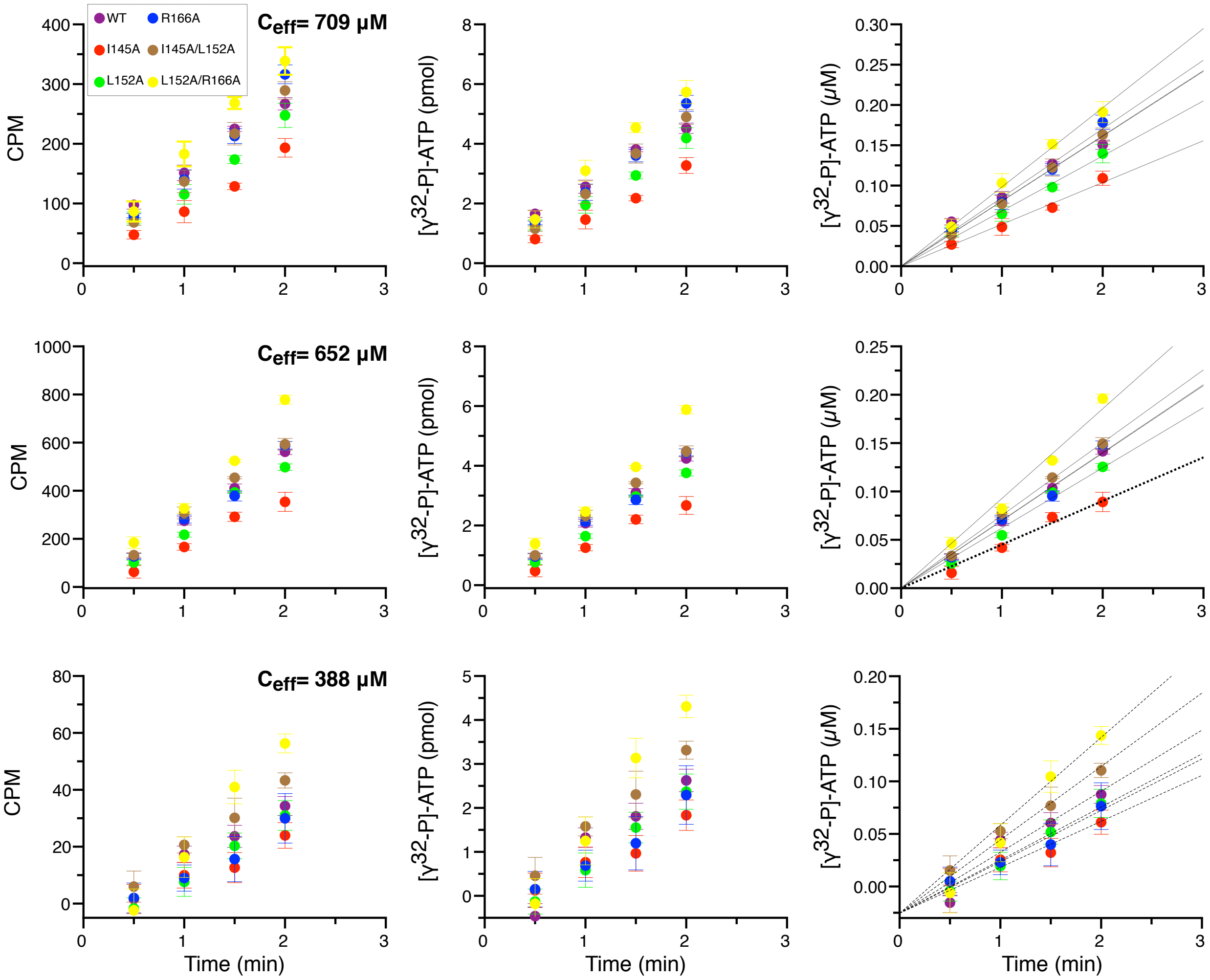


**Figure S9: Measurement of steady-state phosphorylation rates in vitro of p66α-QS-WT substrate for three different effective concentracions.** These measurements were performed at 3 µM substrate concentration and using as enzyme constructs: MBD2-(GS)$_{10}$-PKAc ($C_{eff}$ =709 µM), MBD2-(GS)$_{30}$-PKAc ($C_{eff}$ =652 µM) and MBD2-(GS)$_{60}$-PKAc ($C_{eff}$ =388 µM). Corrected counts per minute (CPM) from $^{32}$P-incorporated to substrate, obtained at different time-steps for each docking interaction variant. Each point was measured by triplicate (Left plot). Pmol of [γ$^{32}$P]-ATP incorporated per minute, obtained after dividing CPM by the slope of the [γ$^{32}$P]-ATP standard curve (Middle plot). Initial velocities were obtained from the slope of the linear regression of µM of $^{32}$P-incorporated to substrate per minute (first-order conditions). Error bars represent the standard error of the mean, with n= 3.

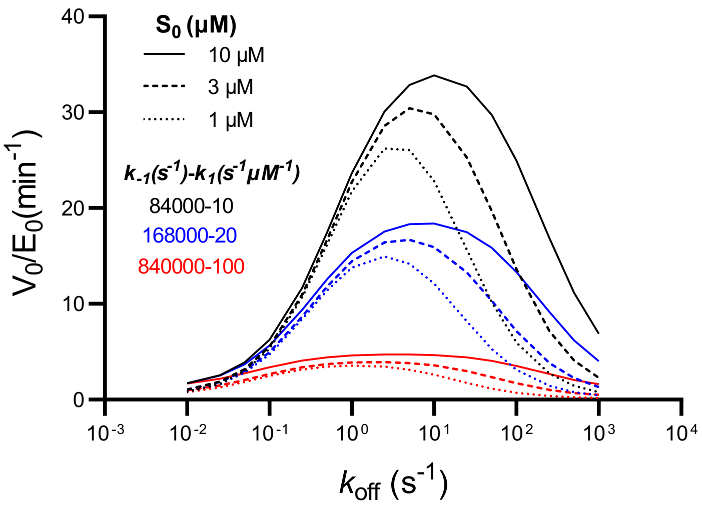


**Figure S10: Steady-state phosphorylation rate depends on substrate association/dissociation rate constants (*$k_1$, $k_{-1}$*)** Variation in phosphorylation rate of R-3K substrate using three different pairs of association-dissociation rate constants. For all cases the ratio $k_{-1}/k_1$ remains constant. As predicted by *Eq. I*, at a given concentration, an n-fold increase of the association-dissociation rate constants results in an n-fold decrease of the phosphorylation rate.

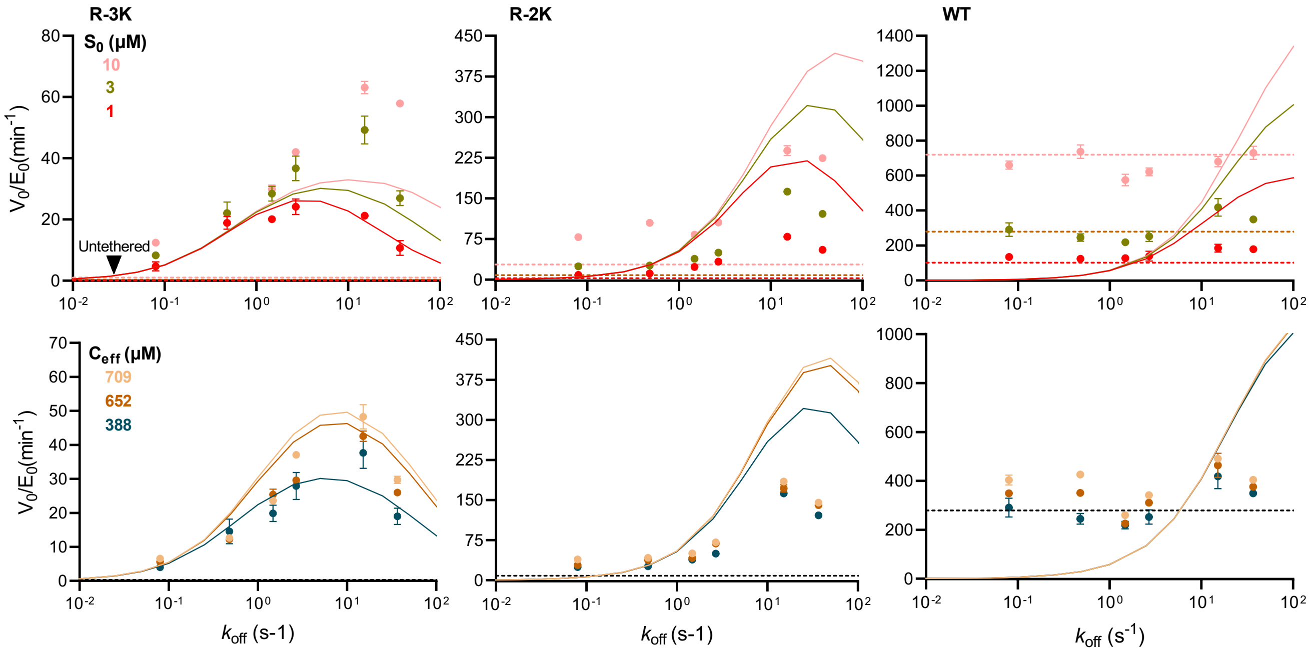


**Figure S11: Steady state phosphorylation rates obtained from experiments and predicted from analytical solutions.** Experimental rate values and the standard error of the mean, with n= 3 (solid circles). *Eq. I* was calculated using a $k_{on}$ value fixed at $1.8 \cdot 10^7$ s$^{-1}$ M$^{-1}$ for a series of $k_{off}$ varying from 0.01–100 s$^{-1}$ (solid line). The dashed lines depict the simulated rates from the untethered reactions with identical parameters and the solid lines represent predicted rates from a reaction that only proceeds through the tethered route based on *Eq. I*.

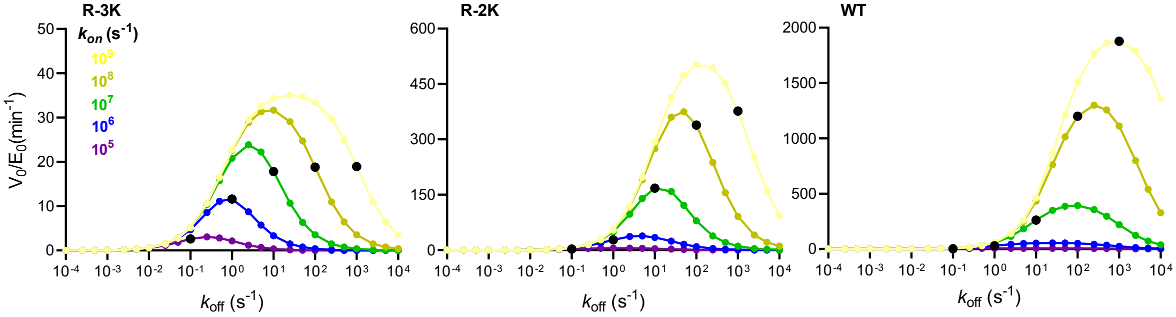


**Figure S12: Phosphorylation rate dependency on $k_{on}$.** Steady-state rate predicted by *Eq. I* for the tethered reaction path at a substrate concentration of 1 μM. Docking interactions with $K_D$ of 1 μM are highlighted as black circles. Parameters used for calculation of *Eq. I* are identical to those in Fig 3.

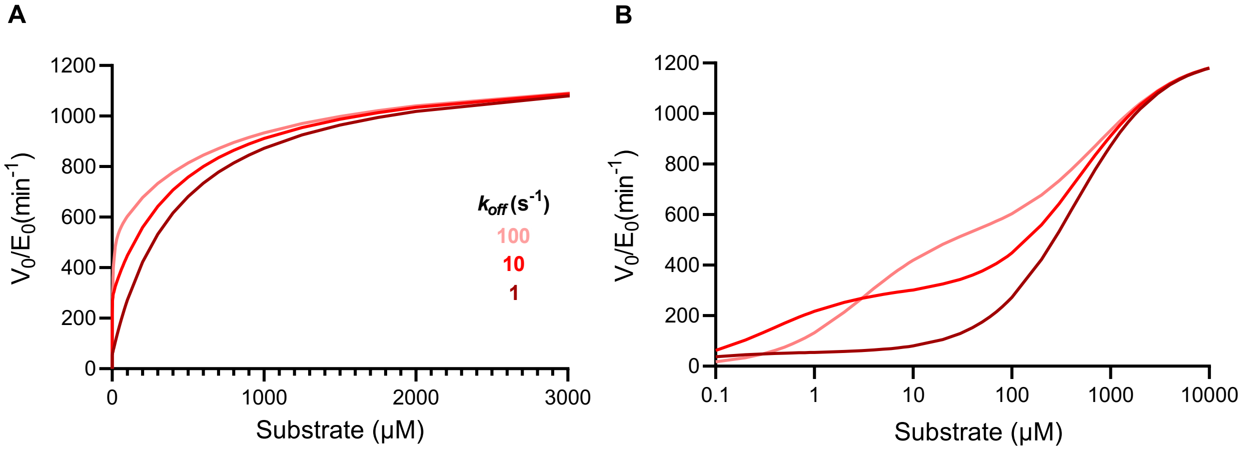


**Figure S13: Phosphorylation rate enhancement dependency on substrate concentration**. Values were predicted by Numerical Simulations using the same parameters as in Fig. 3, with substrate concentration represented in linear (A) and logarithmic scale (B). The rate enhancement for the proposed model seems to follow a Michaelis-Menten-like dependency at a narrow range of substrate concentrations (A), however is clearly not Michaelis-Menten-like in the entire concentration range (B). The enhancement of docking interactions at affinities below the apex is analogous to the decrease in $K_M$ ($k_{off}$ = 100 $s^{-1}$), and the decrease at high affinities is analoguous to decreasing $k_{cat}$ ($k_{off}$ = 1 $s^{-1}$).

#### Supplementary tables:

**Table S1: Rate constants from stopped-flow experiments.**

| p66α variant | $k_{on}$ ($s^{-1}\mu M^{-1}$) ± S.E. | $k_{off}$ ($s^{-1}$) ± S.E. | $K_D$ (μM) ± S.E. | $K_D/K_D(WT)$ |
|---|---|---|---|---|
| WT | 18.2 ± 0.30 | 0.0772 ± 0.0002 | 0.0042 ± 0.00007 | 1 |
| R166A | 14.6 ± 0.86 | 0.483 ± 0.0015 | 0.033 ± 0.002 | 7.8 |
| L152A | 10.5 ± 0.21 | 2.69 ± 0.017 | 0.256 ± 0.005 | 61 |
| I145A | 8.99 ± 0.46 | 1.48 ± 0.027 | 0.165 ± 0.009 | 39 |
| L152A/R166A | 8.78 ± 0.60 | 15.2 ± 0.11 | 1.7 ± 0.12 | $4.0 \times 10^2$ |
| I145A/L152A | 3.70 ± 0.025 | 36.8 ± 0.76 | 9.9 ± 0.2 | $2.4 \times 10^3$ |

#### Derivation of an analytical equation describing the rate of phosphorylation under reversible tethering

**Strategy:**

The goal is to express the rate of product formation as a function of elemental rate constants and total concentrations of enzyme and substrate. The full reaction scheme can be studied by numerical simulations, but is too complex for an analytical solution. We thus make simplifying assumptions that allow us to deduce an analytical solution for the curve shape describing the dependence of the tethered reaction on the docking parameters. We also predict the docking strength that produces the highest phosphorylation rate.

**Definitions:**

We study an idealized tethered system as defined in (Dyla & Kjaergaard (2020) *PNAS* 117 (35) 21413-21419). Briefly, the system consists of a kinase tethered to its substrate via a heterodimeric coiled-coil docking interaction. The kinase and the substrate motif are linked to each part of the docking interaction via a flexible linker of variable length. In total, the linkers define the effective concentration of intra-complex substrate binding, $C_{eff}$. The tethering is reversible and defined by the rate constants $k_{on}$ and $k_{off.}$ (Figure 1B). Binding of the substrate to the active site of the kinase is governed by the rate constants $k_1$ and $k_{-1}$. The rate constant for irreversible phospho-transfer is denoted $k_2$ and the rate constant for product dissociation, $k_3$.

**Assumptions:**

We wish to derive the initial steady state phosphorylation rate, which describes time scales much longer than the slowest reaction steps, but sufficiently short to assume that the substrate concentration is unchanged and the phosphorylated substrate can be ignored after it has dissociated from the docking interaction. We therefore assume:

$[P] = 0, [S] = S_0$

Furthermore, we aim to study scenarios where tethered phosphorylation is dominant, which is generally true for low substrate concentrations. Mechanistically, this is the equivalent of assuming that the ternary complexes (OSS, OSP, OPS, OPP, ES and EP shown in Figure 1B and S1) are unimportant for the overall phosphorylation rates. We therefore derive the equation based on the 'Tethered' kinetic model shown in Figure 1B. We later validate the appropriateness of these assumptions by comparison of numerical simulation results based on the full kinetic model to the equation derived from the 'Tethered' pathway only. These results were undistinguishable under low substrate concentrations.

**Derivation:**

The Law of Mass Action applied to the 'Tethered' kinetic model (Figure 1B) leads to the following system of nonlinear reaction equations:

$$\frac{dS}{dt} = -k_{on}[E][S] + k_{off}[OS]$$

$$\frac{dE}{dt} = -k_{on}[E][S] + k_{off}[OS] + k_{off}[OP]$$

$$\frac{dOS}{dt} = k_{on}[E][S] - (k_{off} + k_1 C_{eff})[OS] + k_{-1}[CS]$$

$$\frac{dCS}{dt} = k_1 C_{eff}[OS] - (k_{-1} + k_2)[CS]$$

$$\frac{dCP}{dt} = k_2[CS] - k_3[CP]$$

$$\frac{dOP}{dt} = k_3[CP] - k_{off}[OP]$$

$$\frac{dP}{dt} = -k_{on}[E][P] + k_{off}[OP]$$

Quasi-steady-state approximation of all intermediates:

$$\frac{dOS}{dt} = \frac{dCS}{dt} = \frac{dCP}{dt} = \frac{dOP}{dt} = 0$$

$$\frac{dCP}{dt} = 0$$

$$k_2[CS] = k_3[CP]$$

$$[CP] = \frac{k_2}{k_3}[CS] \qquad (1)$$

$$\frac{dOP}{dt} = 0$$

$$k_3[CP] = k_{off}[OP]$$

Inserting (1) into the above equation:

$$k_3 \cdot \frac{k_2}{k_3}[CS] = k_{off}[OP]$$

$$[OP] = \frac{k_2}{k_{off}}[CS] \qquad (2)$$

$$\frac{dCS}{dt} = 0$$

$$k_1 C_{eff}[OS] = (k_{-1} + k_2)[CS]$$

$$[CS] = \frac{k_1 C_{eff}}{k_{-1} + k_2}[OS] \qquad (3)$$

$$\frac{dOS}{dt} = 0$$

$$k_{on}[E][S] - \left(k_{off} + k_1C_{eff}\right)[OS] + k_{-1}[CS] = 0$$

Inserting (3) into the above equation:

$$k_{on}[E][S] - \left(k_{off} + k_1C_{eff}\right)[OS] + \frac{k_{-1}k_1C_{eff}}{k_{-1} + k_2}[OS] = 0$$

$$k_{on}[E][S] = (k_{off} + k_1C_{eff} - \frac{k_{-1}k_1C_{eff}}{k_{-1} + k_2})[OS]$$

$$[OS] = \frac{k_{on}[E][S](k_{-1} + k_2)}{(k_{off} + k_1C_{eff})(k_{-1} + k_2) - k_{-1}k_1C_{eff}} = \frac{k_{on}[E][S](k_{-1} + k_2)}{k_{off}k_{-1} + k_{off}k_2 + k_1k_2C_{eff}}$$

$$[OS] = \frac{k_{on}[E][S]}{k_{off} + \frac{k_1k_2C_{eff}}{k_{-1} + k_2}} \qquad (4)$$

We are interested in the rate of product [P] formation:

$$\frac{dP}{dt} = -k_{on}[E][P] + k_{off}[OP]$$

Inserting (2) into the above equation:

$$\frac{dP}{dt} = -k_{on}[E][P] + k_{off} \cdot \frac{k_2}{k_{off}}[CS]$$

Inserting (3) into the above equation:

$$\frac{dP}{dt} = -k_{on}[E][P] + k_2 \cdot \frac{k_1C_{eff}}{k_{-1} + k_2}[OS]$$

Inserting (4) into the above equation:

$$\frac{dP}{dt} = -k_{on}[E][P] + \frac{k_1k_2C_{eff}}{k_{-1} + k_2} \cdot \frac{k_{on}[E][S]}{k_{off} + \frac{k_1k_2C_{eff}}{k_{-1} + k_2}}$$

$$\frac{dP}{dt} = -k_{on}[E][P] + \frac{k_{on}k_1k_2C_{eff}[E][S]}{k_{off}k_{-1} + k_{off}k_2 + k_1k_2C_{eff}} \qquad (5)$$

From the conservation law for the enzyme, total enzyme concentration is constant:

$$E_0 = [E] + [OS] + [CS] + [CP] + [OP]$$

$$[E] = E_0 - ([OS] + [CS] + [CP] + [OP])$$

Inserting (1), (2), and (3) into the above equation:

$$[E] = E_0 - ([OS] + \frac{k_1C_{eff}}{k_{-1} + k_2}[OS] + \frac{k_2}{k_3} \cdot \frac{k_1C_{eff}}{k_{-1} + k_2}[OS] + \frac{k_2}{k_{off}} \cdot \frac{k_1C_{eff}}{k_{-1} + k_2}[OS])$$

$$[E] = E_0 - (1 + \frac{k_1 C_{eff}}{k_{-1} + k_2} + \frac{k_1 k_2 C_{eff}}{k_3(k_{-1} + k_2)} + \frac{k_1 k_2 C_{eff}}{k_{off}(k_{-1} + k_2)})[OS]$$

$$[E] = E_0 - \frac{k_{off} k_3 (k_{-1} + k_2) + k_{off} k_1 k_3 C_{eff} + k_{off} k_1 k_2 C_{eff} + k_1 k_2 k_3 C_{eff}}{k_{off} k_3 (k_{-1} + k_2)}[OS]$$

$$[E] = E_0 - \frac{k_{off} k_3 (k_{-1} + k_2 + k_1 C_{eff} + \frac{k_1 k_2 C_{eff}}{k_3} + \frac{k_1 k_2 C_{eff}}{k_{off}})}{k_{off} k_3 (k_{-1} + k_2)}[OS]$$

Inserting (4) into the above equation:

$$[E] = E_0 - \frac{k_{-1} + k_2 + k_1 C_{eff} + \frac{k_1 k_2 C_{eff}}{k_3} + \frac{k_1 k_2 C_{eff}}{k_{off}}}{k_{-1} + k_2} \cdot \frac{k_{on}[E][S]}{k_{off} + \frac{k_1 k_2 C_{eff}}{k_{-1} + k_2}}$$

$$[E] = E_0 - \frac{(k_{-1} + k_2 + k_1 C_{eff} + \frac{k_1 k_2 C_{eff}}{k_3} + \frac{k_1 k_2 C_{eff}}{k_{off}})k_{on}[S]}{k_{off} k_{-1} + k_{off} k_2 + k_1 k_2 C_{eff}}[E]$$

$$[E](1 + \frac{\left(k_{-1} + k_2 + k_1 C_{eff} + \frac{k_1 k_2 C_{eff}}{k_3} + \frac{k_1 k_2 C_{eff}}{k_{off}}\right)k_{on}[S]}{k_{off} k_{-1} + k_{off} k_2 + k_1 k_2 C_{eff}}) = E_0$$

$$[E] = \frac{(k_{off} k_{-1} + k_{off} k_2 + k_1 k_2 C_{eff})E_0}{k_{off} k_{-1} + k_{off} k_2 + k_1 k_2 C_{eff} + \left(k_{-1} + k_2 + k_1 C_{eff} + \frac{k_1 k_2 C_{eff}}{k_3} + \frac{k_1 k_2 C_{eff}}{k_{off}}\right)k_{on}[S]}$$

Substituting [E] into the product formation equation (5):

$$\frac{dP}{dt} = -k_{on}[E][P] + \frac{k_{on} k_1 k_2 C_{eff}[S]}{k_{off} k_{-1} + k_{off} k_2 + k_1 k_2 C_{eff}} \cdot \frac{(k_{off} k_{-1} + k_{off} k_2 + k_1 k_2 C_{eff})E_0}{k_{off} k_{-1} + k_{off} k_2 + k_1 k_2 C_{eff} + \left(k_{-1} + k_2 + k_1 C_{eff} + \frac{k_1 k_2 C_{eff}}{k_3} + \frac{k_1 k_2 C_{eff}}{k_{off}}\right)k_{on}[S]}$$

Initial rates: $[P] = 0, [S] = S_0$

$$\frac{dP}{dt} = \frac{k_{on} k_1 k_2 C_{eff} E_0 S_0}{k_{off} k_{-1} + k_{off} k_2 + k_1 k_2 C_{eff} + \left(k_{-1} + k_2 + k_1 C_{eff} + \frac{k_1 k_2 C_{eff}}{k_3} + \frac{k_1 k_2 C_{eff}}{k_{off}}\right)k_{on} S_0}$$

$$\frac{dP}{dt} = \frac{k_1 k_2 C_{eff} E_0 S_0}{(k_{-1} + k_2 + \frac{k_1 k_2 C_{eff}}{k_{off}})\frac{k_{off}}{k_{on}} + \left(k_{-1} + k_2 + k_1 C_{eff} + \frac{k_1 k_2 C_{eff}}{k_3} + \frac{k_1 k_2 C_{eff}}{k_{off}}\right)S_0} \quad (6)$$

Equation (6) is grouped based on $k_{off}$:

$$\frac{dP}{dt} = \frac{k_1 k_2 C_{eff} E_0 S_0}{(\frac{k_{-1} + k_2}{k_{on}}) k_{off} + k_1 k_2 C_{eff} S_0 \frac{1}{k_{off}} + \left(k_{-1} + k_2 + k_1 C_{eff} + \frac{k_1 k_2 C_{eff}}{k_3}\right) S_0 + \frac{k_1 k_2 C_{eff}}{k_{on}}}$$

Substitute parameters of the equation above to make the functional form more clear:

$$a = k_1 k_2 C_{eff} E_0 S_0$$

$$b = \frac{k_{-1} + k_2}{k_{on}}$$

$$c = k_1 k_2 C_{eff} S_0$$

$$d = \left(k_{-1} + k_2 + k_1 C_{eff} + \frac{k_1 k_2 C_{eff}}{k_3}\right) S_0 + \frac{k_1 k_2 C_{eff}}{k_{on}}$$

$$\frac{dP}{dt} = \frac{a}{b k_{off} + c \frac{1}{k_{off}} + d} \qquad (I)$$

#### Prediction of optimal tethering strength.

**Strategy:** Optimal tethering strength is achieved for such $k_{off}$ value, at which product formation rate $\left(\frac{dP}{dt}\right)$ reaches maximum. Thus, to find the apex of the volcano plot, we need to find the maximum of equation (6) with respect to $k_{off}$, and assuming constant $k_{on}$.

We derive (6) relative to $k_{off}$ by applying the chain rule:

$$\frac{d}{dk_{off}} \left[ \frac{a}{b k_{off} + c \frac{1}{k_{off}} + d} \right] = a \cdot \frac{d}{dk_{off}} \left[ \frac{1}{b k_{off} + c \frac{1}{k_{off}} + d} \right] = -a \cdot \frac{\frac{d}{dk_{off}} \left[ b k_{off} + c \frac{1}{k_{off}} + d \right]}{\left( b k_{off} + \frac{c}{k_{off}} + d \right)^2}$$

$$= -\frac{a \left( b - \frac{c}{k_{off}{}^2} \right)}{\left( b k_{off} + \frac{c}{k_{off}} + d \right)^2}$$

The maximum of $\frac{dP}{dt}$ is found at derivative equal to zero, when $k_{off} = k_{off\_apex}$:

$$-\frac{a\left(b-\frac{c}{{k_{off\_apex}}^2}\right)}{\left(bk_{off\_apex}+\frac{c}{k_{off\_apex}}+d\right)^2}=0$$

$$b-\frac{c}{{k_{off\_apex}}^2}=0$$

$$\frac{c}{{k_{off\_apex}}^2}=b$$

$${k_{off\_apex}}^2=\frac{c}{b}$$

$$k_{off\_apex}=\sqrt{\frac{c}{b}}$$

Replace substituted variables back with rate constants:

$$k_{off\_apex}=\sqrt{\frac{k_{on}k_1k_2C_{eff}S_0}{k_{-1}+k_2}} \qquad (II)$$

For enzymes that follow classical Michaelis-Menten kinetics, we can use the following simplifications:

Given $K_{\mathrm{M}}=\frac{k_{-1}+k_2}{k_1}$ and $k_{cat}=k_2$:

$$k_{off\_apex}=\sqrt{\frac{k_{cat}}{K_{\mathrm{M}}}k_{on}C_{eff}S_0} \qquad (III)$$
